# Robust Chromosome Segmentation in Sparse Single-Cell Whole-Genome Sequencing Using a k-Segment Constrained Hidden Markov Model

**DOI:** 10.64898/2026.10.19.760240

**Authors:** Trevor Ridgley, Peter Chovanec, Shiyang He, Yi Yin

**Affiliations:** Department of Human Genetics, David Geffen School of Medicine, UCLA, Los Angeles, CA, 90095, USA

## Abstract

Genome integrity is essential for cellular and organismal fitness. Loss of heterozygosity (LOH), which eliminates the buffering effect of heterozygosity and can expose deleterious recessive alleles, is an important form of somatic genome alteration associated with aging and tumorigenesis. To characterize LOH across cell populations, we use sci-L3, a high-throughput combinatorial-indexing approach for single-cell whole-genome sequencing (WGS). The throughput, however, comes at the cost of sparse coverage of individual genomes, complicating LOH inference from phased heterozygous sites. We therefore developed a *k*-segment constrained hidden Markov model (HMM) that explicitly limits the maximum number of state transitions and implicitly enforces learning a much lower transition probability than unconstrained HMM models, enabling robust segmentation with learned transition probabilities and optional SNP binning. Benchmarking against orthogonal sci-L3-Strand-seq segmentation, which resolves all seven mitotic crossover outcomes, the method achieved 86% LOH recall and 88% precision. Haplotype-aware analyses support genuine LOH in 68-78% of sampled apparent false-positive calls, suggesting that the reference-based precision of 88% may underestimate the actual precision, potentially approaching 96%. The *k*-segment HMM thus enables sensitive detection of LOH heterogeneity from sparse single-cell WGS and is applicable to systems with available haplotype phasing.

## Introduction

### Loss of heterozygosity (LOH) and single-cell detection

LOH was first demonstrated by Curt Stern in 1936 through his discovery of somatic recombination producing reciprocal homozygous “twin spots” in *Drosophila* (1), and now a recognized driver of genome alteration in cancer (2). LOH inactivates tumor suppressors including RB1 in retinoblastoma, osteosarcoma, and non-small-cell lung cancer (3, 4) and BRCA1/BRCA2 and other homologous-recombination (HR) genes in breast and ovarian cancer (5–7). Recently, population-scale studies have linked large copy-neutral LOH (cnLOH) hematopoietic cells to clonal expansion and hematologic-malignancy risk (8–10). These studies have sufficient power to associate both rare variants and polygenic risk scores derived from common variants with the direction of clonal expansion. They further show that cnLOH is a common mosaic chromosomal alteration in older individuals and that most detected events are large, often encompassing substantial portions of chromosome arms. However, these analyses rely on SNP arrays and bulk whole-genome sequencing (WGS), in which population-level allelic imbalance preferentially reveals clonally expanded events. Consequently, their observed prevalence conflates the rate at which structural variants (SVs) arise with the subsequent selection of cells carrying them. Single-cell sequencing can separate these processes by detecting SVs as they arise, before clonal expansion shapes their representation. Although scaling to large populations forces sparse per-cell coverage, large chromosomal alterations spanning many megabases (Mb) remain detectable. Measuring these events directly across individual cells can therefore reveal their underlying frequencies in a polyclonal population.

Mechanistically, large-scale LOH can arise through three major processes: end joining of a chromosome double strand break (DSB) (11); reciprocal mitotic crossover; and 3) nonreciprocal break-induced replication (BIR). End-joining of a single DSB typically produces small deletions of 1 to 50 bp, whereas large interstitial or terminal deletions require two coincident DSBs, which are uncommon in the absence of exogenous DNA damage (**Fig.1A**). HR initiated by a single lesion, in contrast, can generate large-scale LOH through nonallelic HR (NAHR) between ectopic repeats, mitotic crossover (**Fig.1B**), or BIR (**Fig.1C**), extending from the initiating lesion to the chromosome end and encompassing tens of Mb. Crossover and BIR produce similar terminal LOH patterns when only one descendant cell is examined because these outcomes are reciprocal in the two daughter cells for crossover but not for BIR, motivating single-cell approaches that can resolve the two mechanisms.

**Figure 1:**
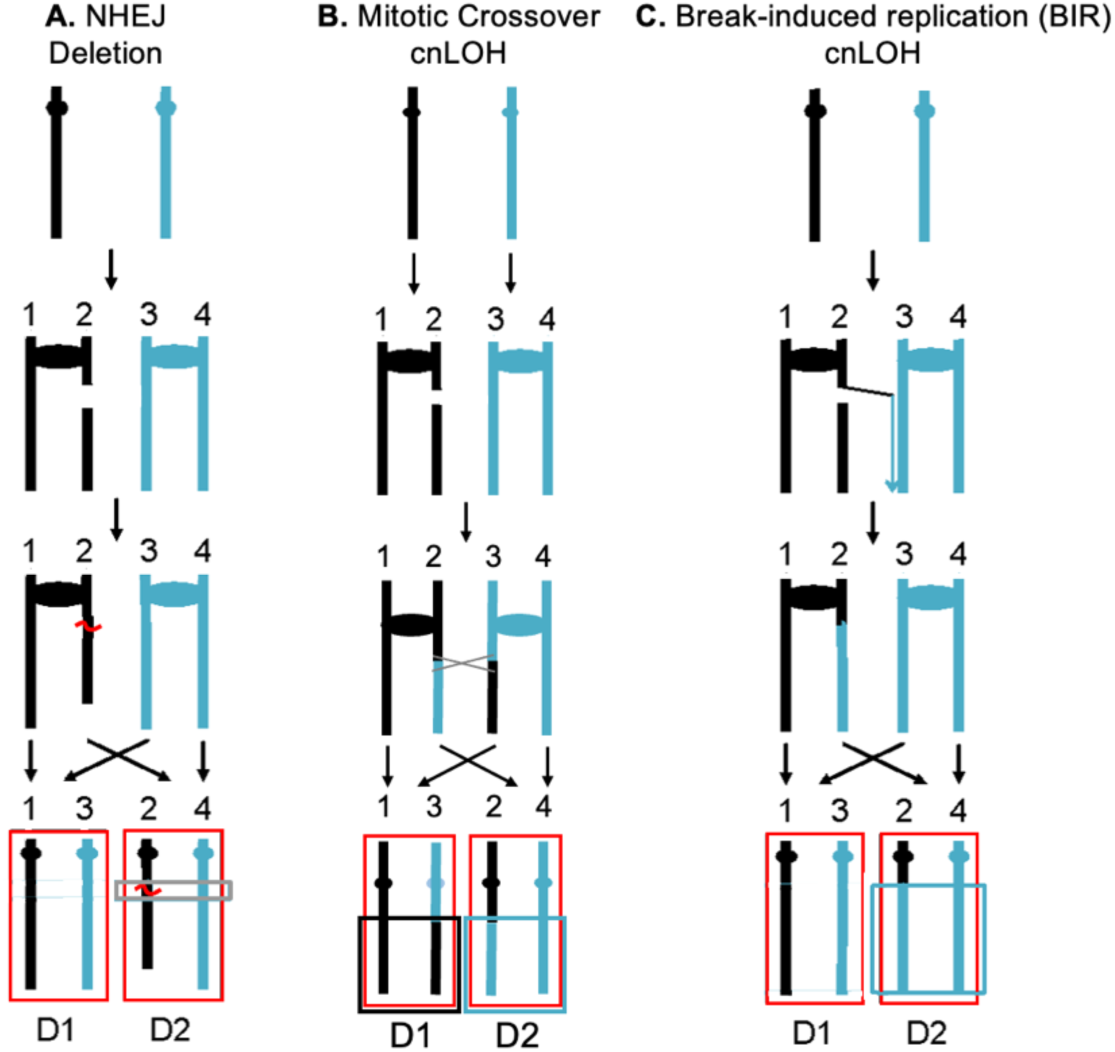
Loss of heterozygosity (LOH) mechanisms. Black and blue depict chromosome homologs. We show DSB in one of the replicated sister chromatids. All three repair mechanisms can cause LOH. **A.** In NHEJ, two broken DNA ends are ligated resulting in an interstitial deletion of the intervening sequences (shortened black chromosome homolog in one of the daughter cells). **B.** Repair of a single DSB by mitotic crossover can cause reciprocal large-scale terminal cnLOH: both homologs are black centromere-distal to the DSB in one cell and blue in the other. The allele frequency between black and blue is still 2:2 without subclonal expansion. Single-cell sequencing can bypass the need to isolate daughter cell clones for analyzing reciprocal cnLOH. **C.** BIR results in cnLOH in one of the two daughter cells. The imbalanced haplotype can be detectable by bulk sequencing, which may or may not reflect subclonal expansion.

### Sparse single-cell WGS and k-segment constrained HMMs

Advances in single-cell genomics have increased throughput while reducing amplification bias. High-coverage methods (∼90% per cell) detect large-scale LOH readily but at high cost (12–14), whereas scalable methods trade coverage per cell for the ability to profile larger cell populations (15–17). We build on sci-L3, a 3-round combinatorial-indexing method coupled with linear amplification for low-cost, high-throughput single-cell WGS (17). Its ∼1% per-cell coverage suits detection of rare genome-instability events across large cell populations and subclonal structure resolution, but creates a statistical challenge: individual heterozygous SNPs are rarely sampled more than once, precluding direct observation of heterozygosity at most loci. LOH must therefore be inferred from the sparse distribution of haplotype-assigned reads along each chromosome. *Yin et al.* previously used three-state HMMs to model haplotype transitions in 1C spermatids and crossover-associated LOH in 2C spermatocytes. Despite the rarity and transient nature of 2C spermatocytes after meiosis I, the throughput of sci-L3 enabled crossover mapping at single-cell resolution and revealed whole-genome equational chromosome segregation during mammalian meiosis I. These results illustrate the power of sparse, high-throughput single-cell WGS for resolving rare chromosome-scale events, while also highlighting the need for robust segmentation methods tailored to sparse genomic observations.

Conventional HMMs have important limitations for sparse single-cell WGS: multinomial models can over-segment, Gaussian models require coverage-dependent binning, and fixed parameters require extensive tuning. Three-state models also cannot readily distinguish LOH from trisomy, generating false-positive (FP) LOH calls that require post-processing. These issues are compounded by large coverage variation: even amplification bias-minimized sci-L3 varies ∼10-fold in coverage within an experiment, and coverage spans orders of magnitude across methods and organisms (15, 17), *e.g.*, sci-L3 samples ∼20%, ∼1%, and ∼0.1% of the genome per cell in yeast, mammalian WGS, and its ATAC adaptation, respectively (18). We therefore adapted the *k*-segment constrained HMM framework of Titsias *et al.* (2016) (19), in which a hyperparameter k bounds the maximum number of state transitions while the number of hidden states is specified independently. Titsias *et al.* developed this framework for copy-number segmentation of bulk WGS, with k controlling resolution. Here, we extend it to sparse single-cell WGS and haplotype-based emissions for LOH detection, to our knowledge, its first application to either setting.

Crossover-associated LOH events are typically large, often spanning much of a chromosome arm, and rare; more than one event per chromosome arm is seldom observed in mitotic cells from yeast to humans, spontaneously or after exogenous damage (8–10, 20, 21). Meiotic crossovers are likewise tightly regulated by interference, suppressing closely spaced events (22). A low k-segment constraint (≤5 segments) is therefore biologically appropriate: it permits the strongest state transitions while suppressing spurious segmentation, whereas increasing k improves model fit at the cost of additional FP LOH calls. A central obstacle to developing and evaluating single-cell LOH callers, however, is the lack of ground truth: clonality can corroborate LOH or CNV calls, but this assumes FP calls occur randomly across cells, an assumption that need not hold.

### An orthogonal strand switch-based chromosome segmentation benchmark

Our lab recently developed sci-L3-Strand-seq (18, 23), which combines the throughput of sci-L3 with Strand-seq to selectively sequence parental DNA template strands. Changes in Watson (W) and Crick (C) strand inheritance provide a direct means of segmenting chromosomes and identifying structural rearrangements, independent of sequence coverage and haplotype emissions (23). For example, deletion-associated LOH converts the normal WC state to W or C, while cnLOH can be distinguished by a single haplotype inheriting both W and C. Template-strand directionality also enables highly accurate genome-wide phasing. In sparse sci-L3-WGS, identifying where to test for LOH is fundamentally a chromosome segmentation problem. Thus, sci-L3-Strand-seq provides an orthogonal benchmark for evaluating HMM-based segmentation: one segments chromosomes by strand inheritance and the other by sparse haplotype observations. These segments, together with phased heterozygous SNPs, provide the inputs for LOH calling.

To evaluate chromosome segmentation into heterozygous and LOH states, we analyzed 931 single cells from the Patski cell line using sci-L3-Strand-seq. Changes in W-C strand orientation provide an orthogonal means of chromosome segmentation, after which phased haplotype fractions can be calculated within each segment. Heterozygous regions contain reads from both parental haplotypes at an expected ratio of approximately 1:1, whereas LOH regions contain predominantly one haplotype. Integrating haplotype fractions with read depth and strand inheritance therefore enables classification of distinct SV outcomes. Across 931 cells, we identified 27 segmental LOH events, 302 uniparental disomies (UPDs), 1,297 monosomies, and 769 segmental losses/deletions. Together, these data provide an orthogonal segmentation benchmark for developing and evaluating the *k*-segment HMM and demonstrate its potential to recover large-scale LOH directly from sparse, phased single-cell WGS.

## Results

### Detecting LOH without Strand-seq segmentation using conventional HMMs

In the development of sci-L3-Strand-seq, we detected chromosomal rearrangements in 931 Patski single cells (23). LOH calling used 28.2 million phased SNPs across the mm10 genome and chromosome segments defined independently by template strand switch. In **Fig.2A**, for example, transitions from CC to W and then WC define three segments on chromosome 9. At a median genome coverage of 0.54% (**Fig.2B**), most heterozygous sites are represented by zero or one read, making direct observation of both alleles at individual sites rare (**Fig.S1A**). Nevertheless, large chromosome segments typically contain more than 400 informative phased SNPs even at ∼1% genome coverage (**Fig.S1B**), providing sufficient information to distinguish heterozygous regions, which contain a mixture of both haplotypes, from LOH regions dominated by one haplotype. We therefore computed haplotype fractions within each strand switch-defined segment, with values near 0 or 1 indicating LOH. Together with copy number and genomic location, these patterns distinguish segmental loss, monosomy, cnLOH, and uniparental disomy (UPD) (**Fig.2C**). Up to five segments were observed per chromosome arm (**Fig.2D**). Because segment boundaries are defined independently by template-strand orientation, these data provide an orthogonally segmented reference for evaluating LOH detection from phased SNPs alone. However, sci-L3-Strand-seq requires cells to undergo at least one round of DNA replication in the presence of BrdU labeling, limiting its application to samples amenable to cell culture or *in vivo* labeling. Many patient-derived specimens, including formalin-fixed paraffin-embedded samples, are incompatible with these requirements. In contrast, sci-L3-WGS and other single-cell WGS methods do not require strand labeling but lack the intrinsic chromosome segmentation provided by Strand-seq, motivating a computational approach to infer LOH segments directly from sparse, phased single-cell WGS.

**Figure 2.**
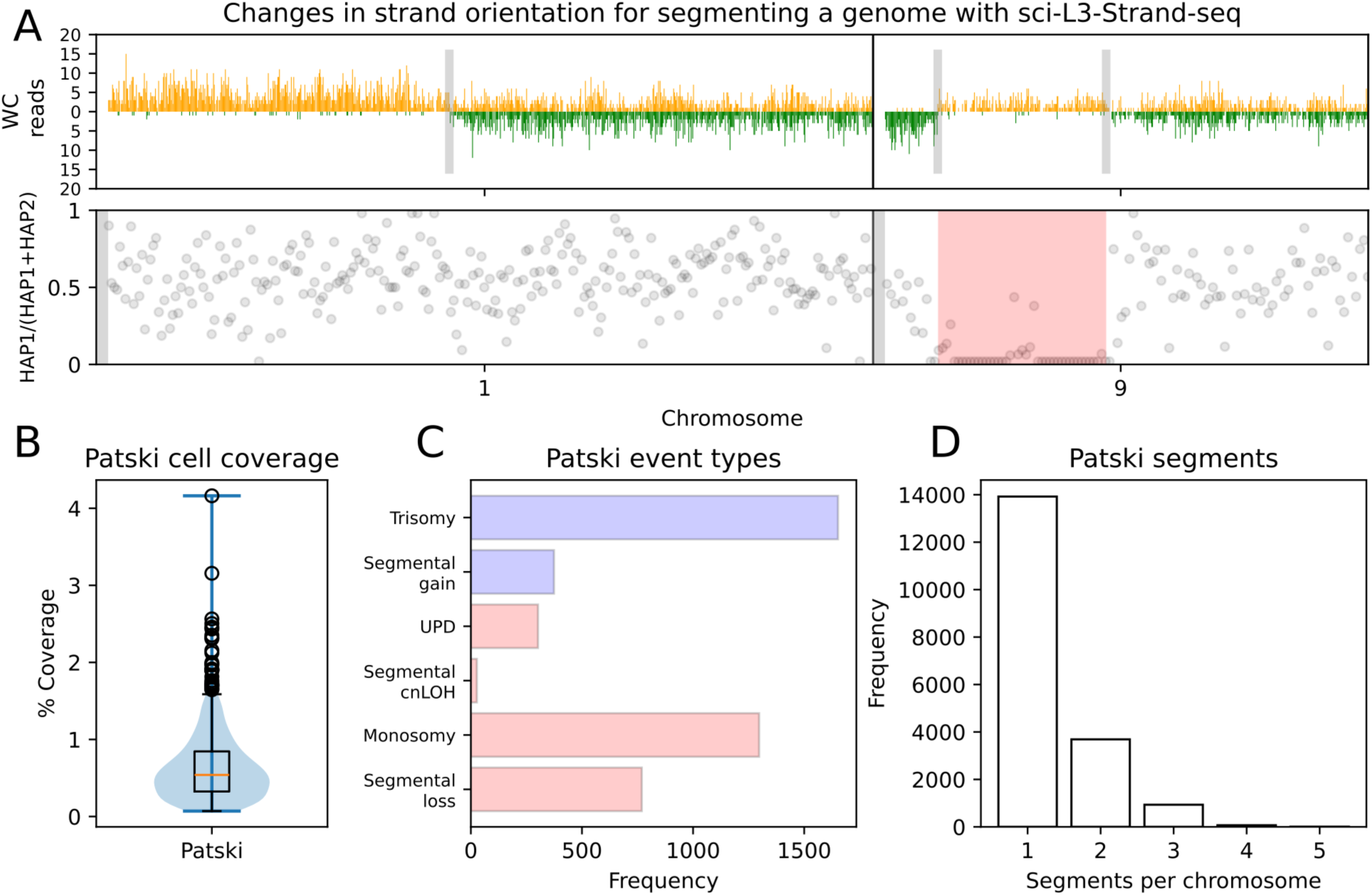
Sci-L3-Strand-seq Patski examples, coverage, annotation and segment numbers. **A.** Two chromosomes from a representative Patski cell. Upper: reads per 200-kb bin colored by template strand directionality (W, yellow; C, green); gray lines indicate changes in strand state. Chr1 transitions from WW to WC, consistent with a sister chromatid exchange (SCE). Chr9 contains three segments (CC-W-WC); the middle segment is single-copy and exhibits LOH (W rather than WW), consistent with segmental loss. Lower: haplotype fraction calculated in bins of ∼40 phased SNPs. Chr1 remains heterozygous across the SCE, whereas the middle segment of chr9 has a haplotype fraction near 0, confirming loss of one haplotype. **B.** Distribution of per-cell genome coverage, with a median of 0.54% (∼20 reads/Mb). **C.** Structural variations identified by sci-L3-Strand-seq. Red denotes LOH events, including segmental loss and cnLOH and whole-chromosome monosomy and UPD; blue represents copy-number gains, including segmental gain and trisomy. **D.** Number of segments per chromosome after post-processing. Only seven chromosome arms contained five segments; therefore, the maximum number of segments (k) for the HMM was set to four.

Without strand information, LOH must be inferred from the phased haplotype sequence alone. We benchmarked three-state Multinomial and Gaussian HMMs for LOH detection, previously used to identify meiotic crossovers from binary haplotype sequences in 1C and 2C cells, respectively (17). For both models, start probabilities were uniform and transition probabilities were fixed at 1 × 10^-10^ divided by the number of SNPs (Multinomial) or bins (Gaussian) per chromosome. The heterozygous-state emission was fixed at 0.5, whereas the two LOH-state emissions were set to 0.9/0.1 for the Multinomial model and 0.95/0.05 for the Gaussian model.

To qualitatively compare LOH calling using sci-L3-Strand-seq segmentation and fixed-parameter HMMs, we first examined a cell sequenced at median coverage that contains several types of LOH. Sci-L3-Strand-seq identified whole-chromosome LOH on chromosomes 4 and 12, an interstitial deletion on chromosome 9, and terminal LOH on chromosome 14 (**Fig.3A**). A standard Multinomial HMM with learned emission and transition probabilities recovers most of the LOH regions on chromosomes 4 and 9, but produces a fragmented call on chromosome 12 and detects only a small portion of the terminal LOH on chromosome 14 (**Fig.3B**). Fixing the transition probabilities substantially reduces this fragmentation: the sci-L3 Multinomial HMM, using the low transition probability defined above, recovers all four LOH events without FP calls (**Fig.3C**).

**Figure 3.**
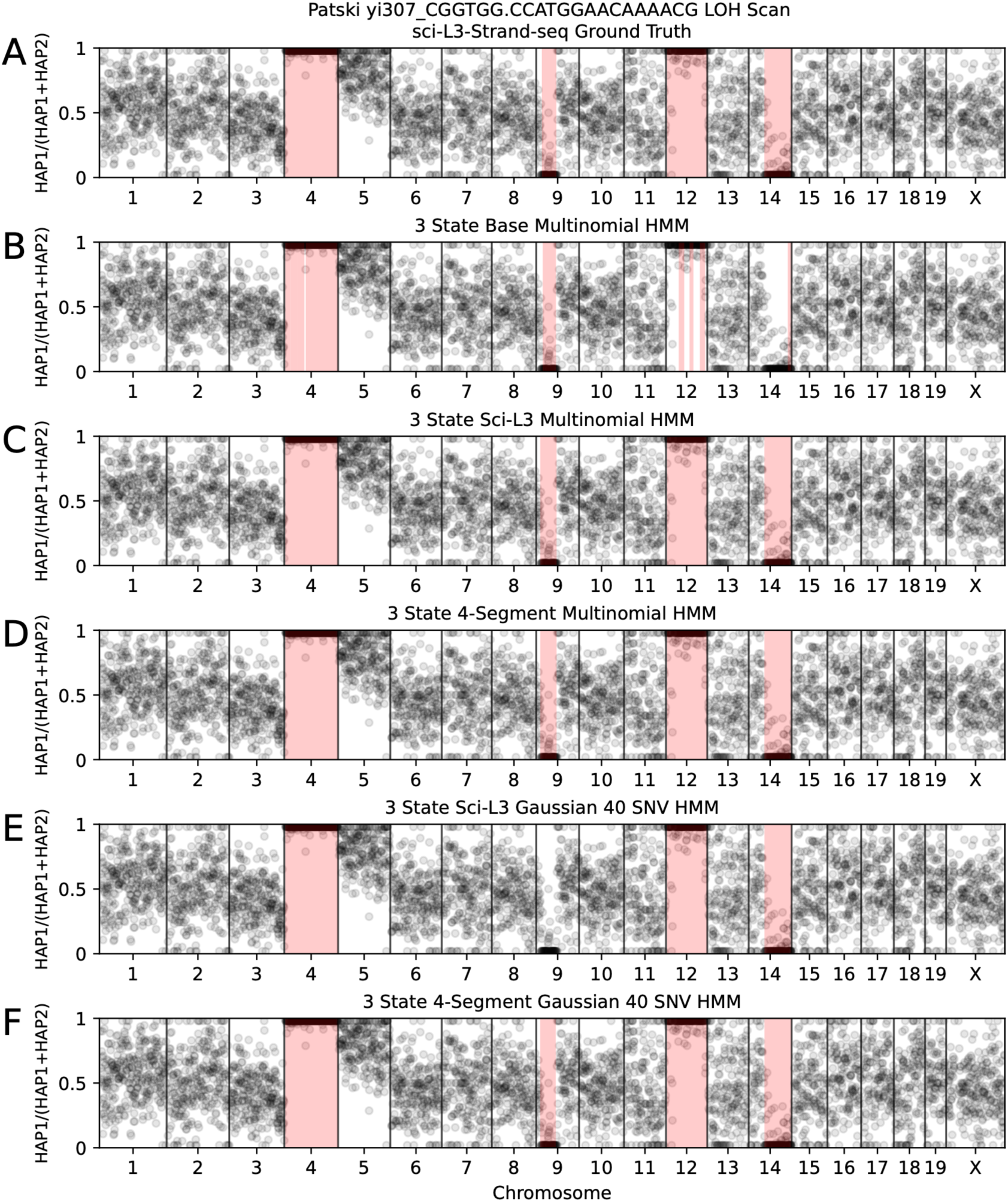
LOH calling in a representative median-coverage Patski cell. Points represent binned haplotype fractions and LOH regions shaded in red. **A.** Sci-L3-Strand-seq reference LOH calls, with Whole-chromosome LOH occurs on chromosomes 4 and 12, interstitial LOH on chromosome 9, and terminal LOH on chromosome 14. Sci-L3-Strand-seq-defined LOH regions serve as the reference (“ground truth”) for comparison with HMM-based calls in panels **B-F**. **B.** LOH calls from a three-state Multinomial HMM with learned emission and transition probabilities. The unfiltered model produces thousands of short segments (27,294 segments; **Tab.S1**), each containing few phased SNPs. After applying 10-Mb size and 20-SNP evidence filters followed by LOH merging, seven calls remain; these include fragmented or incomplete recovery of the reference LOH regions on chromosome 12 and detection of only a small portion of the terminal LOH on chromosome 14. **C.** LOH calls from the three-state sci-L3 Multinomial HMM, which uses fixed emission and transition probabilities to limit over-segmentation. In this median-coverage cell, all four reference LOH events are recovered without FP calls. **D.** LOH calls from the three-state, four-segment (k=4) Multinomial HMM with the same fixed emissions as in **C** but learned transition probabilities under the k-segment constraint. The model recovers both whole-chromosome LOH events, the interstitial LOH on chromosome 9, and the terminal LOH on chromosome 14. **E.** LOH calls from the three-state sci-L3 Gaussian HMM using binned haplotype fractions and fixed parameters. Three of the four reference LOH events are recovered, while the interstitial LOH on chromosome 9 is missed. **F.** LOH calls from the three-state, four-segment Gaussian HMM with learned transition probabilities under the k-segment constraint (k=4). The model recovers all four reference LOH events, including the interstitial LOH on chromosome 9 missed by the fixed-parameter Gaussian HMM in **E**.

Single-cell sequencing noise and the uneven genomic distribution of informative SNPs cause haplotype observations to deviate from the simple binomial expectation of independent sampling (e.g., a single read may be the only read covering multiple adjacent SNPs, all from one haplotype). Consequently, a standard Multinomial HMM can overfit such noise by learning excessively high transition probabilities, producing thousands of short LOH segments in the absence of additional size or SNP evidence filters. Even adjacent regions with modest differences in haplotype fraction (e.g., 0.90 versus 0.95) can introduce additional state transitions and contribute to fragmented calls. Fixing the transition probability avoids this overfitting but introduces a different limitation: although the heuristic used by the sci-L3 Multinomial HMM performs well in the median-coverage example, it does not generalize across the wide range of single-cell sequencing depths and performs particularly poorly, and somewhat counterintuitively, in high-coverage cells (**Figs.S3-S5; Table S1**).

### LOH detection using a k-segment HMM in a representative Patski cell

To address these challenges, we implemented a k-segment constrained HMM, in which k limits the number of segments in the underlying chromosome state path. We set k = 4 primarily based on the expected biology of mitotic recombination, allowing a chromosome arm to contain both a segmental and a terminal LOH event. This choice was also supported empirically by sci-L3-Strand-seq, where chromosomes with more than four segments were rare (**Fig.2D**). The k-segment constraint was implemented in the hmmlearn Python package for both Multinomial and Gaussian HMMs. Here, k specifies the maximum number of segments (one plus the number of state transitions) per chromosome arm, while allowing the most probable state path to contain ≤ *k* segments. At each step *t* along the observation sequence, state probabilities are updated while separately tracking paths containing 1 through *k* segments, thereby preventing any inferred path from exceeding k segments (**Fig.S2**; *Methods*). By limiting the number of state transitions, the k-segment constraint prevents the HMM from over-segmenting noisy haplotype observations into numerous short states, even in regions with haplotype fractions near 0 or 1 (**Fig.S3B**, unfiltered). Applied to the Multinomial HMM with fixed emission parameters, the *k*-segment constraint eliminates the fragmented LOH calls in the example cell while recovering the Strand-seq-defined LOH regions (**Fig.3D**).

An alternative strategy for reducing overfitting to sequencing noise is to bin the binary haplotype observations and model the resulting haplotype fractions (0 to 1) with a Gaussian HMM. Binning reduces the influence of individual noisy observations and is expected to reduce FP LOH calls, but at the cost of sensitivity to shorter segmental events. Consistent with this trade-off, the fixed-parameter sci-L3 Gaussian HMM recovers three of the four LOH events in the example cell but misses the interstitial deletion on chromosome 9 (**Fig.3E**). Applying the k-segment constraint allows transition probabilities to be learned while controlling over-segmentation and recovers the missed chromosome 9 event (**Fig.3F**).

The effect of the k-segment constraint is also apparent in the number of inferred chromosome segments. The sci-L3-Strand-seq reference contains 25 segments across the 20 chromosomes of the median-coverage example cell. In comparison, the standard three-state Multinomial HMM with fixed emissions produces 9,944 segments (**Tab.S1, Fig.S3-5**), reflecting severe over-segmentation. The four-segment Multinomial HMM reduces this number to 74 while allowing transition probabilities to be learned. The fixed-parameter Gaussian HMM produces only 21 segments, but this reduced segmentation comes at the cost of sensitivity, as illustrated by the missed interstitial LOH on chromosome 9 (**Fig.2C, 3E**). Importantly, segmentation by the k-segment HMM remains controlled at higher sequencing coverage. In a second Patski cell with 3.16% genome coverage and 30 sci-L3-Strand-seq-defined segments (**Fig.S5**), the standard Multinomial HMM with fixed emission probabilities but learned transitions produces 44,536 segments, whereas the sci-L3 Multinomial HMM with fixed, low transition probabilities still produces 341 segments. In contrast, the four-segment Multinomial HMM produces 74 segments. The fixed-parameter Gaussian HMM produces 26 segments but again tends to sacrifice sensitivity to segmental LOH. Thus, constraining segmentation provides a means to control over-segmentation across substantially different sequencing depths without relying on extremely low fixed transition probabilities.

### Controlling over-segmentation across the single-cell library

We next examined chromosome segmentation by each HMM across the entire single-cell library. Multinomial HMMs generally produce more segments than Gaussian HMMs, consistent with modeling individual rather than binned SNP observations (**Fig.4A**). Conventional Multinomial HMMs show substantial over-segmentation, with outlier chromosomes containing hundreds to thousands of segments. The k-segment constraint largely eliminates these extreme outliers, although the median number of segments remains higher than for the Gaussian models. To determine whether this additional segmentation represents LOH, we excluded heterozygous segments and, for five-state models, copy-number gains (**Fig.4B**). Conventional Multinomial models retain numerous outlier LOH segments, whereas the k-segment Multinomial HMM shows a substantially lower median. Most of its additional segmentation in **Fig.4A** instead reflects adjacent heterozygous segments with slightly different learned emission probabilities (e.g., 0.53 and 0.48 rather than 0.5 across the chromosome arm) and therefore does not generate additional LOH calls. Consequently, the number of LOH segments produced by the k-segment Multinomial HMM approaches that of the binned Gaussian models, suggesting effective control of over-segmentation.

**Figure 4.**
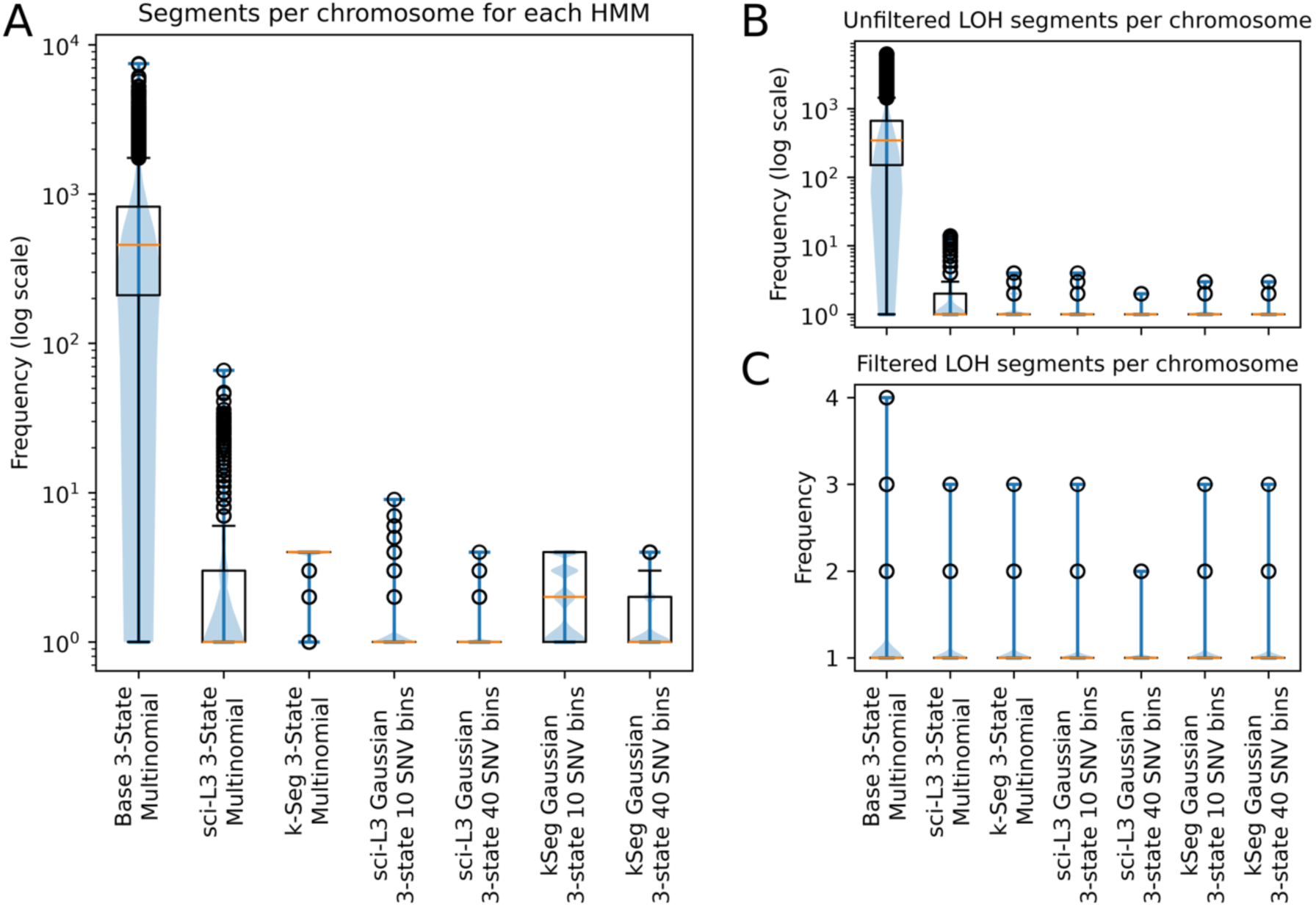
Comparison of chromosome segmentation and LOH calls across HMMs. Each boxplot shows the distribution of segment counts across all 20 chromosome arms in 931 Patski cells for seven HMMs. Red horizontal lines indicate medians, and circles indicate outliers beyond 1.5x the interquartile range. **A.** Total raw chromosome segments, including heterozygous, LOH, and gain states. **B.** Unfiltered LOH segments per chromosome after excluding heterozygous and gain states. **C.** LOH segments per chromosome after applying size and SNP evidence filters.

Next, we examined whether outlier LOH segments could be removed by size (10 Mb) and SNP evidence filters (20 SNPs, **Fig.4C**). Filtering substantially reduces the number of LOH segments produced by Multinomial models, but has little effect on Gaussian HMMs, which already have a median of one LOH segment per chromosome and narrow interquartile ranges. The k-segment HMMs similarly exhibit few segmentation outliers, although with slightly broader interquartile ranges than the Gaussian models, a distinction examined further in the LOH sensitivity analysis below.

### Benchmarking LOH event detection against strand-switch-defined, orthogonal sci-L3-Strand-seq segmentation

To assess overall LOH calling performance, we defined LOH events in the sci-L3-Strand-seq reference set as deletions, monosomies, cnLOH, or UPD. An HMM call was considered a true positive (TP) if the inferred LOH state overlapped at least 60% of the reference LOH event by genomic length; otherwise, the event was considered a false negative (FN). This event-based metric avoids weighting performance by the number of informative heterozygous SNPs within each event, which would favor larger events containing more SNPs. Thus, each reference LOH event contributes equally to performance assessment, regardless of its genomic size or SNP evidence. The number of reference segments and events in each class is summarized in **Tab.S2**.

We first benchmarked LOH detection across seven three-state HMMs with fixed emission probabilities. Raw LOH segments were defined by haplotype fractions ≤0.1 or ≥0.9 and subsequently required to span at least 10 Mb and contain at least 20 informative SNPs (hereafter, the “size and evidence filter”; unfiltered calls are shown in **Fig.S3**). After filtering, the standard Multinomial HMM with unconstrained learned transition probabilities detected 1,915 of 2,395 reference LOH events (TPR = 80.0%; **Tab.1**) and incorrectly called LOH in 730 of 22,012 non-LOH reference segments (FPR = 3.32%). The sci-L3 Multinomial HMM with low fixed transition probabilities detected 2,129 LOH events (TPR = 88.9%) with 449 FP calls (FPR = 2.04%). The four-segment Multinomial HMM, which instead learns start and transition probabilities under the segmentation constraint, produced similar recall with somewhat more FP calls, detecting 2,120 LOH events (TPR = 88.5%) with 481 FP calls (FPR = 2.19%). Thus, the k-segment constraint allows transition probabilities to be learned while maintaining LOH detection performance comparable to the empirically tuned fixed-transition model.

While the Multinomial HMM was previously used to call meiotic crossovers in 1C cells, a Gaussian HMM was used for crossover detection in 2C cells after meiosis I (17). Applied to the 931 Patski cells, the sci-L3 Gaussian HMM with 40-SNP bins produces 305 fewer FP LOH calls than the sci-L3 Multinomial HMM (144 *vs.* 449), reducing the FPR from 2.0% to 0.7% (**Tab.1**). This increased stringency comes at a substantial cost in sensitivity, with 372 fewer TP LOH events and a TPR decrease from 88.9% to 73.4%. Reducing the bin size from 40 to 10 SNPs partially restores sensitivity, recovering 210 additional TP events and increasing TPR to 82.1%, but also increases FP calls from 144 to 214.

We next asked whether the k-segment constraint could recover sensitivity lost by the sci-L3 Gaussian HMM while retaining its relatively low FPR. With 40-SNP bins, the four-segment Gaussian HMM detects 283 additional TP LOH events compared with the sci-L3 Gaussian HMM, accompanied by 121 additional FP calls (**Tab.1**). With 10-SNP bins, the k-segment constraint similarly recovers 133 additional TP events, accompanied by 162 additional FPs. Thus, the k-segment constraint substantially recovers sensitivity lost through Gaussian binning, although with a corresponding increase in FP calls. The previously published sci-L3 HMMs were developed for a much smaller dataset of ∼200 2C cells (17), where bin size and transition probabilities could be empirically optimized and candidate LOH calls manually curated. Such parameter tuning and manual curation become impractical for population-scale single-cell datasets, motivating an approach that accommodates variable sequencing coverage with minimal dataset-or cell-specific parameter tuning.

We next compared HMM performance for segmental LOH events (terminal and interstitial) *vs.* whole-chromosome LOH. Whole-chromosome LOH does not require breakpoint inference and can be detected from the chromosome-wide haplotype fraction alone. Accordingly, all seven HMMs detect whole-chromosome LOH with >92% TPR and ≤0.3% FPR (**Tab.S3**). The standard Multinomial HMM has the lowest TPR at 92.7%, whereas the remaining six models achieve ≥96.2%.

Segmental LOH is substantially more challenging to call. The standard Multinomial HMM detects only 432 of 796 reference events (TPR = 54.3%) while producing 697 FP calls (FPR = 3.2%; **Tab.S3**). Sensitivity is lowest for the sci-L3 Gaussian HMM with 40-SNP bins, which detects only 218 segmental LOH (TPR = 27.4%). Introducing the k-segment constraint more than doubles the number detected to 497 (TPR = 62.4%), demonstrating that constrained segmentation can recover much of the sensitivity lost through Gaussian binning. Reducing the bin size from 40 to 10 SNPs further increases TPR from 27.4% to 53.1% for the sci-L3 Gaussian HMM and from 62.4% to 69.2% for the k-segment Gaussian HMM, accompanied by increases in FPR from 0.4% to 0.7% and from 1.0% to 1.5%, respectively. The sci-L3 and k-segment Multinomial HMMs achieve the highest segmental LOH sensitivities (72.7% and 71.7%, respectively), but at higher FPRs of 1.8% and 2.0%. Thus, the principal differences among HMMs emerge in segmental rather than whole-chromosome LOH detection, revealing a tradeoff between sensitivity and FP control.

**Table 1.** Overall LOH calling performance across three-state Hidden Markov Models. All HMMs use three states and were benchmarked against 2,395 LOH and 22,012 non-LOH sci-L3-Strand-seq reference segments from 931 Patski cells. TPs require ≥60% overlap with a reference LOH event; FPs have no overlap threshold and are counted when a filtered LOH call overlaps a reference non-LOH segment. Detailed model parameterization is provided in Tables S3-S5. Derived precision–recall values plotted in Fig.5A. Recall = TP / 2,395 reference LOH; Precision =TP /(TP + FP). Distance from ideal = sqrt((1 - Recall)^2 + (1 - Precision)^2), the distance of each model’s (Recall, Precision) point from the perfect point (1,1).

| Model | Binning | LOH TP | TPR | LOH FP | FPR | Distance |
| --- | --- | --- | --- | --- | --- | --- |
| Standard Multinomial | None | 1,915 | 80.00% | 730 | 3.3% | 0.341 |
| sci-L3 Multinomial | None | 2,129 | 88.90% | 449 | 2.0% | 0.207 |
| k -segment Multinomial | None | 2,120 | 88.50% | 481 | 2.2% | 0.218 |
| sci-L3 Gaussian | 10 SNP | 1,967 | 82.10% | 214 | 1.0% | 0.204 |
| sci-L3 Gaussian | 40 SNP | 1,757 | 73.40% | 144 | 0.7% | 0.277 |
| k -segment Gaussian | 10 SNP | 2,100 | 87.70% | 376 | 1.7% | 0.196 |
| k -segment Gaussian | 40 SNP | 2,040 | 85.20% | 265 | 1.2% | 0.188 |
| sci-L3 Gaussian | Adaptive SNP | 1,963 | 82.00% | 213 | 1.0% | 0.205 |
| k -segment Gaussian | Adaptive SNP | 2,068 | 86.30% | 287 | 1.3% | 0.183 |

Multinomial HMMs achieve higher sensitivity for segmental LOH, whereas Gaussian HMMs better suppress FP calls. To balance these advantages, we implemented adaptive binning in Gaussian HMMs, adjusting bin size according to each cell’s sequencing coverage or number of informative phased SNPs. Relative to coverage-based binning, SNP-based binning recovered slightly more TP LOH events in both the sci-L3 and k-segment Gaussian HMMs; its effect on FP calls was model-dependent, marginally increasing FPs in the sci-L3 Gaussian HMM but reducing them in the k-segment Gaussian HMM (**Tab.S4**). Overall, adaptive binning performed comparably to the best fixed bin sizes, with SNP-based adaptation in the k-segment Gaussian HMM yielding a favorable sensitivity-specificity balance. These observations motivated a precision-recall analysis to identify the optimal HMM configuration.

Finally, we evaluated whether learning emission probabilities could further improve LOH detection. Unlike transition probabilities, which are sensitive to sequencing coverage and SNP density, LOH-state emissions are expected to remain near 0 or 1 because LOH regions contain only one parental haplotype. We therefore compared k-segment Multinomial and Gaussian HMMs with fixed vs. learned emissions and evaluated LOH filtering based on either observed haplotype fractions or learned state emissions (**Tab.1, S5**). With haplotype-fraction filtering, learning emissions increased TP LOH calls by 24 for the Multinomial HMM and 32 for the Gaussian HMM, accompanied by 18 and 20 additional FPs, respectively. Filtering by learned emissions recovered slightly more TP calls (2,150 and 2,110) but added more FPs (505 and 425; **Tab.S5**). Because haplotype-fraction filtering is applied after HMM segmentation, these results suggest that learning emissions may permit additional short or weakly supported LOH segments to be inferred, increasing both true-and false-positive calls. For the Gaussian HMM, learned emissions with haplotype-fraction filtering moved the 40-SNP model closer to ideal (distance 0.181 vs. 0.188 with fixed emissions), a gain comparable to that of adaptive SNP binning with fixed emissions (0.183). We therefore retained fixed emissions for the selected model, which avoids fitting emissions for each chromosome arm from multiple random initializations, while emission-based filtering did not consistently improve LOH detection.

### Precision-recall evaluation of HMMs for LOH detection

To identify the HMM that best balances LOH sensitivity and FP calls, we compared precision and recall across model configurations (**Fig.5A**). The sci-L3-Strand-seq reference contains 2,395 LOH and 22,012 non-LOH segments, making precision particularly informative because even a low FPR can produce substantial numbers of FP calls. We therefore quantified each model’s Euclidean distance from the ideal point of 100% precision and recall. The k-segment Gaussian HMM with adaptive SNP binning showed the best balance among the models in **Tab.1**, with a distance of 0.183 from the ideal, followed closely by the k-segment Gaussian HMM with fixed 40-SNP bins (0.188).

**Figure 5.**
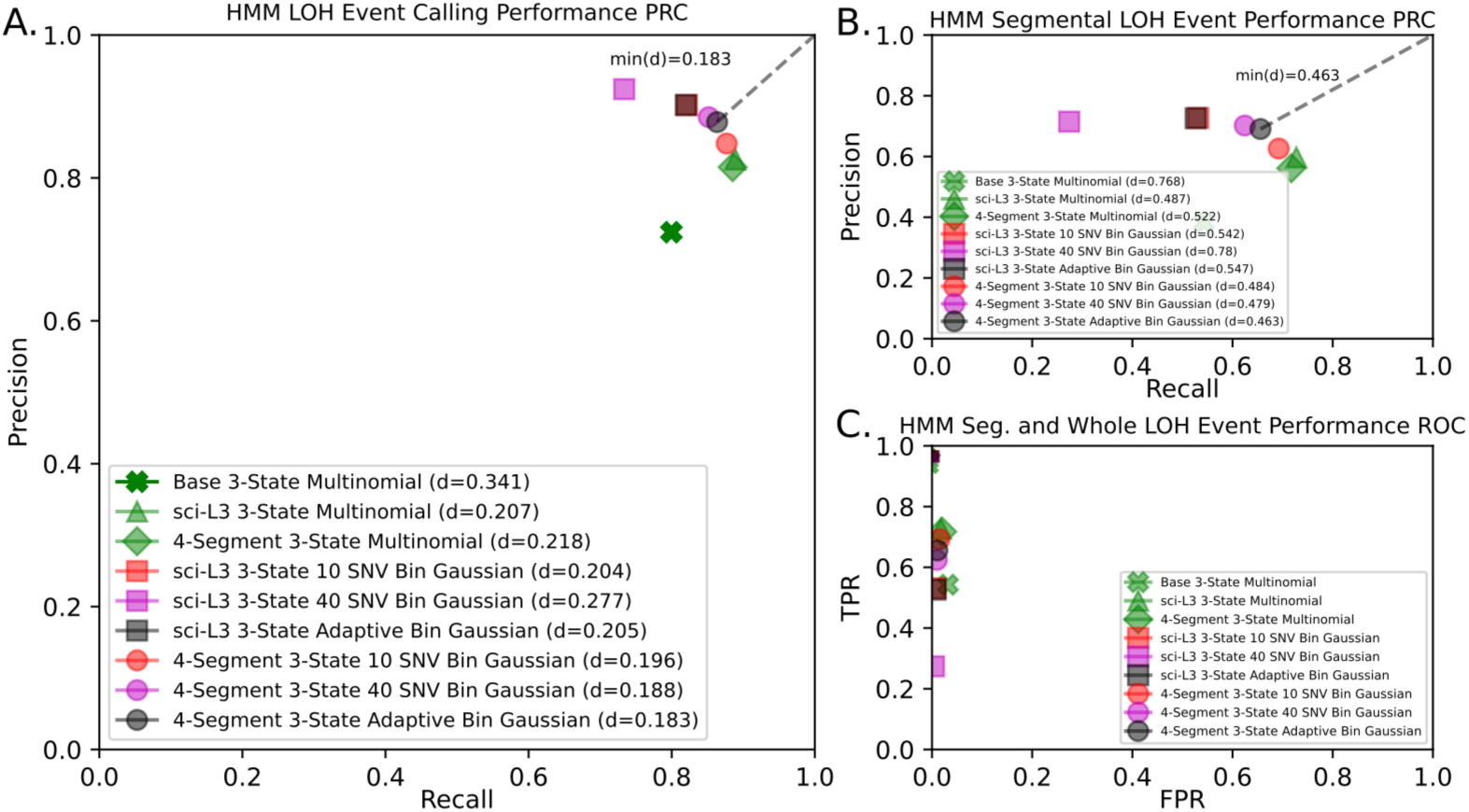
Precision-recall and ROC performance of HMMs for LOH detection. Marker shapes denote HMM categories: standard Multinomial (cross), sci-L3 Multinomial (triangle), k-segment Multinomial (diamond), sci-L3 Gaussian (square), and k-segment Gaussian (circle). Colors indicate binning configurations: 10-SNP bins (red), 40-SNP bins (magenta), adaptive SNP binning (black), and Multinomial models with fixed emissions (green). **A.** Recall (TPR, x-axis) versus precision [TP/(TP+FP), y-axis] across nine HMM configurations. The reference dataset contains 2,395 LOH and 22,012 non-LOH segments. The k-segment Gaussian HMM with adaptive SNP binning is closest to the ideal point (1,1), representing 100% precision and recall. Gaussian HMMs generally achieve higher precision but lower recall than Multinomial HMMs. **B.** Precision-recall performance stratified by segmental LOH (large markers) and whole-chromosome LOH (small markers). Whole-chromosome events cluster near the ideal point (1,1), whereas segmental LOH events more clearly distinguish HMM performance. The k-segment Gaussian HMMs provide the most favorable precision-recall balance for segmental LOH, particularly with adaptive SNP binning. (C) Receiver operating characteristic (ROC) representation of TPR (y-axis) versus FPR (x-axis) across the same nine HMM configurations.

Although Multinomial HMMs avoid binning and the associated loss of information from averaging haplotype observations, this advantage does not translate into superior LOH detection. While Multinomial HMMs perform well in ROC analysis (**Fig.5C**), their precision is lower than that of Gaussian models (**Fig.5A**). For example, the sci-L3 Gaussian HMM with 40-SNP bins produces 144 FPs compared with 449 for the sci-L3 Multinomial HMM (Fisher’s exact test, p=6.8×10^-38^). However, this reduction in FPs comes at a substantial cost in recall, with the 40-SNP sci-L3 Gaussian HMM achieving the highest precision but lowest recall among the models evaluated.

The k-segment constraint substantially recovers recall for both Gaussian bin sizes by allowing transition probabilities to be learned while limiting over-segmentation (40-SNP bins, p=5.1×10^-24^; 10-SNP bins, p=9.3×10^-8^; Fisher’s exact test). Because the number of informative SNPs varies with sequencing coverage, we further evaluated adaptive binning calibrated to the phased SNP count on chromosome 19, the smallest mouse chromosome. Combining adaptive SNP binning with the k-segment constraint produced the most favorable precision-recall balance (**Fig.5A, Tab.1**), with significantly higher recall than the sci-L3 Gaussian HMM using equivalent adaptive binning (p=3.8×10^-5^; Fisher’s exact test).

The distinction between models was more pronounced for segmental LOH (**Fig.5B**). Whole-chromosome LOH was detected with high precision and recall by nearly all models and does not require inference breakpoints. In contrast, segmental LOH requires accurate breakpoint inference and therefore provides a more stringent test of HMM segmentation performance. The k-segment Gaussian HMMs provided the most favorable precision-recall balance for these events, particularly with adaptive SNP binning.

### Haplotype-aware analysis reveals LOH calls missed by sci-L3-Strand-seq segmentation

An important advantage of sci-L3-Strand-seq is its ability to provide chromosome segmentation independently of haplotype-based LOH detection, enabling orthogonal validation of the k-segment HMM. However, strand-switch-based segmentation does not capture all LOH events. First, in a diploid cell, the two parental homologs can inherit four equally likely template-strand configurations: WW, WC, CW, and CC. Segmental loss on WC or CW chromosomes produces a detectable strand switch, whereas the 50% loss on WW or CC chromosomes (e.g., WW to W) may escape strand-based segmentation To rescue these events, we use strand-switch-defined LOH regions from one cell to calculate haplotype fractions in corresponding regions of other cells, identifying shared LOH events without detectable strand switches (**Fig.6A**). However, this clonal rescue may be incomplete. Second, even deletions expected to produce strand switches on WC chromosomes may be missed because of amplification and sequencing noise or aneuploidy that alters the expected strand-state transition (e.g., WWC to WC, **Fig.6B**). Third, cnLOH is by definition copy-neutral and almost never causes strand switching. Both strand-based segmentation and subsequent rescues are typically missed (**Fig.S7**). These limitations suggest that some apparent FP LOH calls by the k-segment HMM may represent genuine events missed by the sci-L3-Strand-seq reference. We therefore used haplotype-aware sci-L3-Strand-seq analysis to independently evaluate these calls (**Figs.6B, S7**).

**Figure 6:**
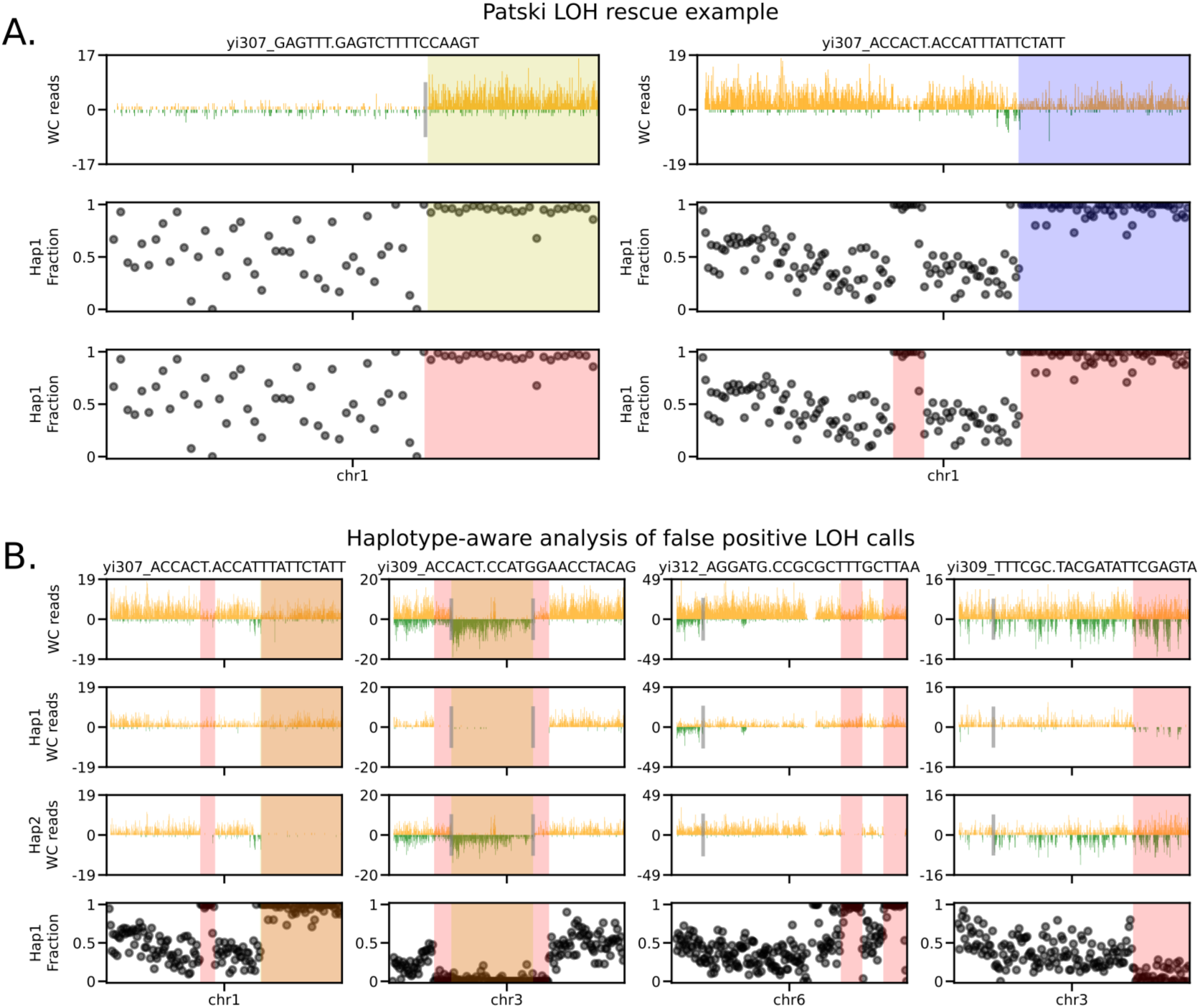
LOH rescue and haplotype-aware analysis of FP LOH calls. **A.** Clonal rescue of an LOH event missed by strand-switch-based segmentation. Left: A strand switch on chromosome 1 (top, gray vertical line) defines a terminal segment with an extreme haplotype fraction, allowing LOH to be called (yellow). Right: A second cell sharing the LOH event lacks a detectable strand switch, causing sci-L3-Strand-seq to treat the entire chromosome arm as a single segment. The resulting chromosome-wide haplotype fraction is not sufficiently extreme to call LOH because the LOH region is averaged with heterozygous regions. Applying the LOH boundary identified in the left cell reveals the shared terminal LOH event in the right cell (blue), which is rescued through clonal analysis. The bottom panels show the corresponding LOH annotations (red) by HMM. **B.** Four examples of apparent FP LOH calls by k-segment HMMs supported by haplotype-aware sci-L3-Strand-seq analysis. For each cell, the top panel shows W and C reads in 200 kb bins; the second and third panels show reads separated by parental haplotype (Hap1 and Hap2), and the bottom panel shows binned Hap1 fractions. HMM LOH calls are highlighted in red, sci-L3-Strand-seq reference calls in yellow, and overlapping regions in orange. 1^st^ col. (chr1; same chr. with **6A** right): The k-segment Multinomial HMM with learned emissions detects a clonally rescued terminal LOH (TP) and an additional interstitial LOH absent from the reference (apparent FP). Haplotype-aware reads support both events by loss of Hap2 reads. 2^nd^ col.(chr3): The sci-L3-Strand-seq reference identifies a central LOH region, whereas the k-segment HMMs detect a larger LOH region extending beyond the reference boundaries (an example of an “LOH-adjacent FP”; WWC to WC transition). Haplotype-aware analysis supports additional loss of Hap1 on both sides of the reference segment, consistent with incomplete segmentation in a partially trisomic chromosome. 3^rd^ col. (chr6): The k-segment HMM identifies an additional segmental LOH region beyond the sci-L3-Strand-seq reference annotation. Haplotype-aware reads support the LOH call in a chromosome with a complex strand configuration. 4^th^ col. (chr3): terminal cnLOH example, sequencing noise and aneuploidy (WWC to WC) complicate strand-switch segmentation. The k-segment HMM detects terminal LOH supported by extreme haplotype fractions and a haplotype-specific WC region.

To perform haplotype-aware segmentation by sci-L3-Strand-seq, reads were first separated by parental haplotype, allowing strand-state changes to be detected independently within each haplotype. For example, a WW to W transition that is invisible to conventional strand-switch segmentation becomes detectable as a loss of reads from one haplotype (“drop-to-zero,” DTZ), while the other remains in the W state. We used these haplotype-specific boundaries to reassess apparent FP LOH calls from the k-segment HMMs. These calls fell into two categories: stand-alone FPs and LOH-adjacent FPs, where HMM-defined LOH extended into neighboring regions annotated as non-LOH by sci-L3-Strand-seq (**Fig.6B** 2^nd^ col., **S7**). We defined LOH-adjacent FPs as extensions of no more than 10 Mb beyond a reference LOH boundary (*e.g.* an HMM call that is slightly larger than a reference LOH).

To estimate the fraction of apparent FP LOH calls that represent genuine events, we randomly sampled 30 stand-alone and 30 LOH-adjacent FP calls from each of three k-segment HMM configurations: Multinomial with learned emissions, Multinomial with fixed emissions, and Gaussian with fixed emissions and adaptive binning. Each event was reexamined using haplotype-aware Strand-seq segmentation to determine whether it could be reclassified as a true LOH event (**Tab.2, S6-11, Fig.S6-S11**).

Among the 30 stand-alone apparent FP LOH calls from the k-segment Multinomial HMM with learned emissions, 21 (70.0%) were supported as genuine LOH events by haplotype-aware analysis (**Tab.2, S6, Fig.S6**). Some represent cnLOH events (e.g., the right end of chromosome 3 in cell 27) that escape strand-switch segmentation because copy-neutral changes do not necessarily alter the WC template-strand configuration (**Fig.6B**). Of the nine remaining FPs, six were associated with drop-to-zero (DTZ) regions, where homozygous deletions or sparse, biased haplotype-specific reads can mimic LOH. Two were associated with copy-number gains, and one lacked sufficient haplotype-aware evidence. These findings suggest that excluding DTZ regions may further reduce FP LOH calls in sparse single-cell WGS.

LOH-adjacent apparent FPs revealed a different limitation of the sci-L3-Strand-seq reference. When LOH events are rescued clonally without detectable strand switches, their boundaries may be imprecisely defined. Consequently, HMM-inferred LOH regions can extend beyond the reference annotation, generating apparent FPs despite strong haplotype evidence (**Fig.6B; Fig.S7; Tab.S7**). Among 30 LOH-adjacent FPs from the learned-emission Multinomial HMM, 26 (86.7%) were supported as genuine LOH, while three were associated with DTZ and one lacked sufficient evidence (**Tab.2**). Thus, HMM-based segmentation can recover more complete LOH boundaries than those inferred through clonal rescue alone.

We next compared apparent FP reclassification across the three k-segment HMM configurations. Haplotype-aware analysis supported 47 of 60 (78.3%) apparent FPs from the learned-emission Multinomial HMM, 45 of 60 (75.0%) from the fixed-emission Multinomial HMM, and 41 of 60 (68.3%) from the Gaussian HMM with adaptive binning (**Tab.2**). Across all models, 133 of 180 (73.9%) apparent FPs showed evidence of genuine LOH, while 32 of the remaining 47 were associated with DTZ regions. The higher reclassification fraction for Multinomial HMMs is consistent with their greater sensitivity to additional LOH events, whereas the Gaussian HMM makes fewer FP calls overall but appears more conservative in detecting LOH.

**Table 2:** Haplotype-aware FP analysis summary across k-Segment HMMs. For each of three k-segment HMM configurations, 30 stand-alone and 30 LOH-adjacent apparent FP calls were randomly sampled and reexamined using haplotype-aware sci-L3-Strand-seq segmentation. LOH-adjacent FPs were defined as HMM calls extending no more than 10 Mb beyond a reference LOH boundary. Calls supported by haplotype-specific evidence were reclassified as likely genuine LOH, whereas the remaining FPs were attributed to drop-to-zero (DTZ) regions, copy-number gains, or HMM segmentation errors/insufficient haplotype-aware evidence. The three configurations differ in emission parameterization and filtering as indicated. Percentages are calculated from 60 sampled calls per model or 180 calls overall.

| Model | Category | Likely real LOH | Total FP | Remain as FP but due to DTZ | Remain as FP but due to gain | FP due to HMM segmentation or lack of HA evidence |
| --- | --- | --- | --- | --- | --- | --- |
| 4-Segment 3-State Multinomial HMM with learned emissions and filtering | Stand-alone FP LOH | 21 | 9 | 6 | 2 | 1 |
|  | LOH-ADJ FP | 26 | 4 | 3 | 0 | 1 |
|  | Total FP LOH | 47 | 13 | 9 | 2 | 2 |
|  | % Total FP LOH | 78.3% | 21.7% | 15.0% | 3.3% | 3.3% |
| 4-Segment 3-State Multinomial HMM with fixed emissions and fraction filtering | Stand-alone FP LOH | 22 | 8 | 4 | 3 | 1 |
|  | LOH-ADJ FP | 23 | 7 | 6 | 0 | 1 |
|  | Total FP LOH | 45 | 15 | 10 | 3 | 2 |
|  | % Total FP LOH | 75.0% | 25.0% | 16.7% | 5.0% | 3.3% |
| 4-Segment 3-State Gaussian HMM with fixed emissions, adaptive binning, and fraction filtering | Stand-alone FP LOH | 17 | 13 | 9 | 3 | 1 |
|  | LOH-ADJ FP | 24 | 6 | 4 | 0 | 2 |
|  | Total FP LOH | 41 | 19 | 13 | 3 | 3 |
|  | % Total FP LOH | 68.3% | 31.7% | 21.7% | 5.0% | 5.0% |
|  | **Total FP LOH** | **133** | **47** | **32** | **8** | **7** |
|  | **% Total FP LOH** | **73.9%** | **26.1%** | **17.8%** | **4.4%** | **3.9%** |

These findings suggest that precision measured against the original sci-L3-Strand-seq reference underestimates the actual precision of k-segment HMMs. For the selected k-segment Gaussian HMM with adaptive SNP binning, the reference-based precision was 87.8%. Haplotype-aware analysis supported genuine LOH in 68.3% of sampled apparent FPs (**Tab.2**). Accounting for likely genuine LOH events among apparent FPs, all three k-segment HMM configurations converge on an estimated precision of approximately 95-96%, suggesting that their apparent differences in precision largely reflect incomplete sci-L3-Strand-seq reference annotation rather than differences in LOH calling accuracy. Thus, beyond its intended application to sparse sci-L3-WGS data, the k-segment HMM can complement strand-switch-based segmentation in sci-L3-Strand-seq by recovering LOH events and refining boundaries missed by the original segmentation.

### Clonal analysis independently validates apparent FP LOH calls

Sparse single-cell WGS is susceptible to stochastic coverage loss, which can produce apparent LOH when one haplotype is underrepresented. However, such sampling artifacts are unlikely to recur with similar boundaries across clonally related cells (14, 24). We therefore used hierarchical clustering of HMM-defined FP LOH profiles to identify recurrent apparent FP calls and assess their clonal support (**Fig.7, S12-13**). We examined three k-segment HMM configurations: Multinomial with learned emissions and emission filtering, Multinomial with fixed emissions and haplotype-fraction filtering, and Gaussian with fixed emissions, adaptive SNP binning, and haplotype-fraction filtering. For the learned-emission four-segment Multinomial HMM, which produced the most apparent FPs (415 segments, compared with 505 FP events in **Tab.S5**), 785 cells clustered together with few or no FP calls (**Fig.7A**). The remaining 87 cells showed recurrent FP patterns, with individual clusters largely confined to single chromosomes (**Fig.7B**), including a few whole-chromosome DTZ events. These can arise when a chromosome that is already monosomic undergoes further loss, potentially through micronucleus formation, leaving very few informative SNPs and generating spurious HMM LOH calls. Because these events reflect chromosome loss rather than conventional LOH, we did not count whole-chromosome DTZ calls as genuine LOH in the clonal FP analysis. We selected chr13 and chrX for detailed analysis because their LOH clusters differed in composition: chr13 clusters consisted almost entirely of reference TP calls, whereas about half of the chrX clusters were dominated by apparent FP calls (**Fig.7C-E**).

**Figure 7.**
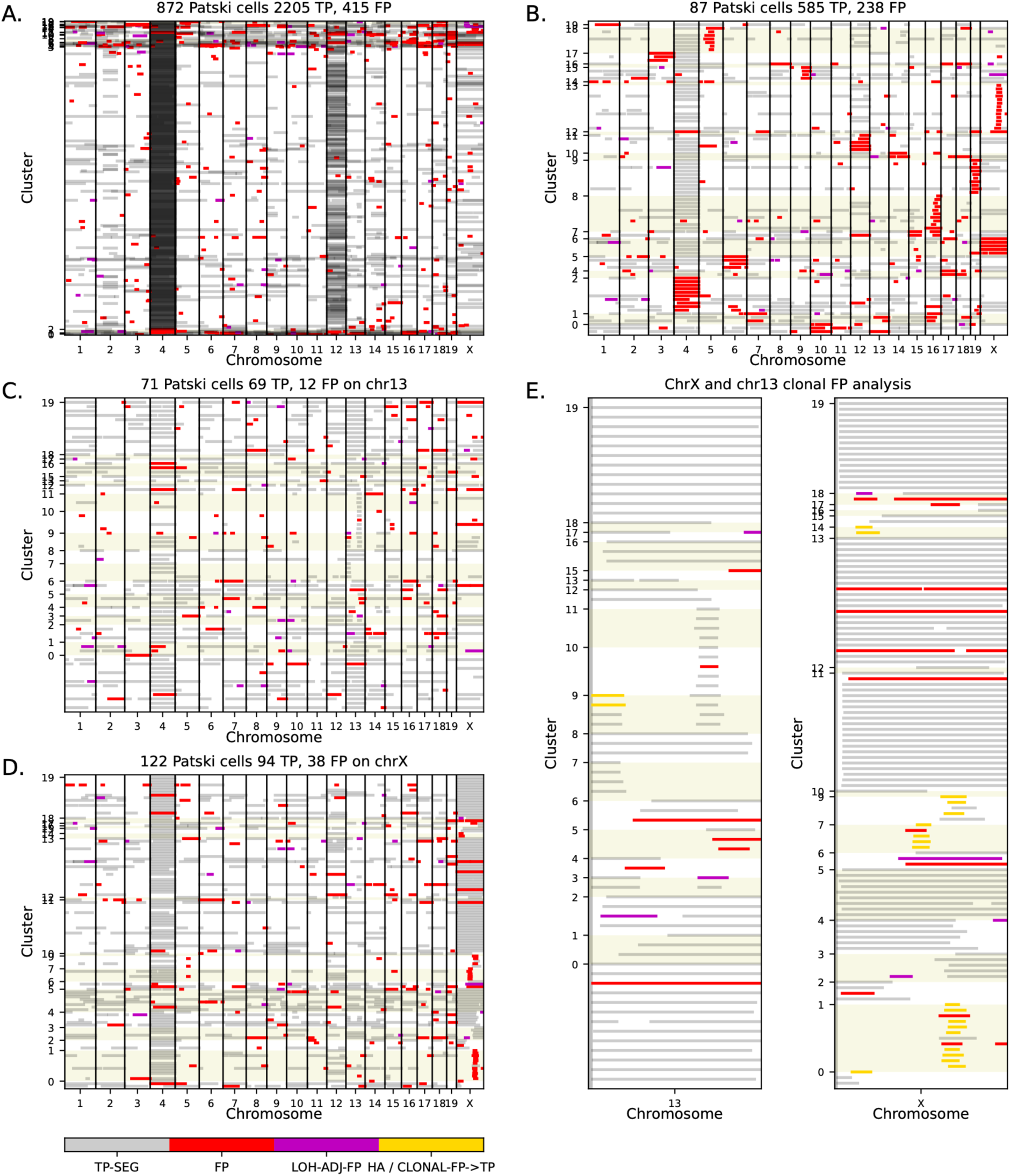
Clustering reveals recurrent apparent FP LOH patterns across Patski cells. LOH calls from the 4-Segment 3-State Multinomial HMM with learned emissions and emission filtering. Each row represents a single cell, with LOH calls encoded by haplotype in 1-Mb genomic bins. Cells were grouped by hierarchical clustering of HMM-defined FP LOH calls. Colors indicate LOH classifications relative to the sci-L3-Strand-seq reference: TP (gray), stand-alone FPs (red), LOH-adjacent FPs (magenta), and apparent FPs reclassified as likely genuine LOH through haplotype-aware or clonal analysis (gold; **E**). Alternating backgrounds distinguish clusters. Counts in panel titles are numbers of HMM LOH segments, rather than projecting them onto the reference LOH calls. **A.** Genome-wide clustering of 872 Patski cells based on apparent FP LOH patterns. Most cells fall into a large cluster with few or no FPs (red or magenta), whereas smaller clusters exhibit recurrent FP patterns. **B.** Genome-wide LOH profiles of 87 cells after excluding the dominant cluster in **A**. Recurrent FP clusters are largely confined to single chromosomes. Clusters with whole-chromosome FPs on chr4 (cluster 2), chr12 (cluster 11) and chrX (cluster 6) are annotated as whole-chromosome DTZ (subsequent loss, likely segregation into micronuclei, from abundant, subclonal monosomy precursors) in the sci-L3-Strand-seq annotations and are not counted as genuine LOH in the clonal analysis. **C.** Genome-wide LOH profiles of 71 cells from chr13-specific clustering. **D.** Genome-wide LOH profiles of 122 cells from chrX-specific clustering. **E.** Enlarged haplotype-specific LOH profiles of chr13 (left) and chrX (right), with cells organized by chromosome-specific clusters. Recurrent apparent FPs were evaluated with haplotype-aware analysis (Tab.S12). Chr13 clusters contain predominantly reference TPs, whereas several chrX clusters contain predominantly apparent FPs, some of which were reclassified as likely genuine LOH (gold) after examining haplotype-aware plots.

Haplotype-aware analysis supported genuine LOH in 19/23 (82.6%) apparent FPs on chrX but 2/6 (33.3%) on chr13. Thus, clusters with few reference TPs could contain many genuine LOH events missed by sci-L3-Strand-seq, whereas the presence of numerous reference TPs did not by itself validate co-clustered FP calls. This suggests that clonal recurrence can identify candidate LOH events, but haplotype-specific evidence remains essential for distinguishing genuine LOH from DTZ regions, copy-number gains, or segmentation errors in the reference. Across the two chromosomes, 21/29 (72.4%) apparent FPs were supported as genuine LOH, broadly consistent with the independent haplotype-aware analysis in **Tab.2**.

We next examined whether these patterns were consistent across HMM configurations (**Fig.S12-13; Tab.S12-14**). The fixed-emission Multinomial HMM produced fewer apparent FP segments than the learned-emission model (389 vs. 415), with 23/41 (56.1%) clonal FPs supported as genuine LOH (**Fig.S12, Tab.S13**), compared with 21/29 (72.4%) for the learned-emission model (Fisher’s exact test, p=0.21; **Tab.S12**). The adaptive-bin Gaussian HMM produced the fewest apparent FP segments (203), and 10/12 (83.3%) of its clonal apparent FPs were supported as genuine LOH (**Fig.S13, Tab.S14**). Thus, the configuration with the fewest apparent FPs also had the highest fraction of them supported as genuine, consistent with more conservative LOH calling in the Gaussian HMM, while LOH events identified by the Multinomial HMMs suggest that these models can recover LOH events missed by the sci-L3-Strand-seq reference. Together, the clonal and haplotype-aware analyses provide complementary evidence that a substantial fraction of apparent FP LOH calls reflect incomplete sci-L3-Strand-seq reference segmentation rather than errors in HMM-based detection.

## Discussion

We adapted the k-segment constrained HMM framework for detecting LOH from sparse, phased single-cell WGS data. The central advantage of this approach is that it incorporates a biologically motivated limit on the number of chromosome segments while allowing transition probabilities to be learned independently for each chromosome arm. Because spontaneous mitotic LOH events are typically large and infrequent, a four-segment constraint per chromosome arm accommodates whole-chromosome, terminal, and interstitial LOH while suppressing excessive state transitions driven by noisy haplotype observations. Unlike conventional HMMs that require empirically fixed transition probabilities to prevent over-segmentation, the k-segment constraint enables parameter learning across cells with widely varying sequencing coverage. In the Multinomial implementation, it also controls over-segmentation without requiring binning of individual SNP observations. Benchmarking against orthogonal sci-L3-Strand-seq segmentation demonstrated that the k-segment HMM achieves robust LOH detection, particularly for segmental events that require accurate breakpoint inference.

The Multinomial and Gaussian implementations reveal an important tradeoff between sensitivity and FP control. Multinomial HMMs retain individual haplotype observations and achieve higher sensitivity for segmental LOH, but also generate more apparent FP calls. Gaussian HMMs suppress noise through binning, reducing FP calls at the expense of sensitivity to shorter LOH events. Introducing the k-segment constraint recovers much of this lost sensitivity by allowing transition probabilities to be learned without excessive segmentation. Adaptive binning based on informative phased SNP counts further accommodates variation in SNP sampling among cells. Among the configurations evaluated, the k-segment Gaussian HMM with adaptive SNP binning provided the most favorable overall precision–recall balance, achieving 86% recall and 88% precision against the sci-L3-Strand-seq reference. Although the improvement over fixed 40-SNP binning was modest, adaptive binning avoids selecting a single bin size for cells with different coverage and SNP densities. Thus, the principal benefit of the k-segment framework is not uniformly higher sensitivity or precision, but robust LOH detection with fewer empirically fixed parameters across heterogeneous single-cell datasets.

An important finding is that even the orthogonal sci-L3-Strand-seq reference does not capture all LOH events even with the clonal rescue scheme. Strand-switch-based segmentation may miss deletions on chromosomes with identical template-strand orientations, cnLOH events that do not alter strand orientation, and changes obscured by aneuploidy or sparse sequencing. Clonal rescue can recover some of these events, but may leave imprecisely defined boundaries. Haplotype-aware analysis supported genuine LOH in 68-78% of sampled apparent FP calls across three k-segment HMM configurations, suggesting that their reference-based precision substantially underestimates their actual precision. Extrapolating these reclassification fractions yields estimated precisions of approximately 95-96% for all three configurations, although these estimates depend on the representativeness of sampled events. Clonal analysis provided complementary evidence by identifying recurrent LOH patterns among apparent FP calls, including events shared across cells but absent from the sci-L3-Strand-seq reference. Notably, clonal recurrence alone did not reliably predict whether an apparent FP was genuine, emphasizing the importance of haplotype-specific validation. These findings suggest that the k-segment HMM can complement strand-switch-based segmentation in sci-L3-Strand-seq by identifying additional LOH events and refining their boundaries, extending its utility beyond the original application to sci-L3-WGS.

### Limitations

Several limitations remain. Our analysis focused on LOH events spanning at least 10 Mb and supported by at least 20 informative phased SNPs, limiting sensitivity to smaller events, particularly in low-coverage cells. Moreover, Patski cells are hybrids between highly diverged mouse strains, with a heterozygous SNP density of approximately 10^-2^, substantially higher than that of typical human genomes. Performance may therefore differ in samples with lower heterozygosity or less complete haplotype phasing. The HMM also identifies LOH states without independently distinguishing cnLOH from deletion or UPD from monosomy, which requires additional copy-number information. Copy-number gains present a related challenge because gain and heterozygous states have overlapping haplotype-fraction distributions, leading to excessive FP gain calls even with constrained segmentation. In addition, whole-chromosome DTZ regions, particularly following further chromosome loss in monosomic cells, can generate misleading LOH calls when very few informative SNPs remain. Integrating read depth, relative chromosome counts, or haplotype-specific coverage could improve LOH subtype classification, gain detection, and DTZ filtering.

## Conclusion

Accurate chromosome segmentation is particularly challenging in sparse single-cell WGS, where LOH must be inferred from limited haplotype observations. To our knowledge, this study provides the first systematic benchmark of single-cell WGS LOH detection against orthogonal, cell-specific chromosome segmentation defined by template-strand switches in sci-L3-Strand-seq. By constraining segmentation while allowing chromosome-arm-specific parameter learning, the k-segment HMM enables robust LOH detection across variable sequencing coverage. Unlike sci-L3-Strand-seq, which requires BrdU labeling and DNA replication, haplotype-based HMM segmentation can be applied to a broader range of biological and clinical specimens, including fixed tissues, as well as studies of whole organisms for which in vivo BrdU labeling is particularly challenging, provided that phased SNP information is available. Conversely, integrating k-segment HMM calls into sci-L3-Strand-seq analysis can recover SVs missed by strand-switch segmentation alone, improving sensitivity to rare chromosome alterations and helping distinguish SCEs from SV-associated strand changes, an essential step toward understanding HR partner choice, which fundamentally determines whether DNA repair preserves genome integrity or promotes genome instability. Together, these complementary approaches expand the ability to detect and characterize spontaneous genome alterations across heterogeneous cell populations.

## Acknowledgments

We thank members of the Yin lab for helpful discussions. This work was supported by the National Institute of Health Training Grant in Genomic Analysis and Interpretation (T32HG002536) to T.R., the National Institute of General Medical Sciences (R35GM142511 to Y.Y), W. M. Keck Foundation (1002444 to Y.Y) and DOD CDMRP (PC250408 to Y.Y.).

## Methods

### Data Pre-Processing

Phased SNPs were assigned to individual cells as described previously for sci-L3-Strand-seq, excluding centromeric and telomeric regions. For Multinomial HMMs, observed haplotypes were encoded as binary sequences (0 or 1), with rare sites covered by both haplotypes represented by adjacent 0 and 1 observations. Sequences were analyzed separately by cell and chromosome arm. For Gaussian HMMs, phased SNPs were grouped into nonoverlapping bins (10, 40, or adaptive SNP counts), with the mean haplotype fraction calculated for each bin.

### k-Segment constrained HMM

LOH states were inferred with HMMs implemented in hmmlearn (Python; development version forked after release 0.2.6, commit 0562ca6). In this version, `MultinomialHMM` emits one categorical symbol per site; in later releases, it was renamed `CategoricalHMM`. `GaussianHMM` was used with diagonal covariance. When unconstrained, these models fit sparse single-cell haplotype sequences with frequent switches between the two homozygous states and largely bypass the heterozygous state. We therefore constrained each hidden path to at most *K* segments (contiguous runs of a single state) per chromosome arm, using the k-segment framework of Titsias et al. (2016). The constraint was applied both during EM parameter learning (the “prospective” use in Titsias et al.) and decoding.

For implementation, we added k-segment versions of hmmlearn’s log-space forward, backward, pairwise-posterior and Viterbi routines (Cython). They are enabled by a single new argument, `smax` = *K*, on `MultinomialHMM` and `GaussianHMM`; when `smax` is unset, the models are unchanged. Titsias et al. use an explicit augmented counting chain. We instead index the forward and backward lattices by segment count:

- *F_c,t_*(*j*) = *P*(*x*_1:*t*_, *z_t_* = *j*, *c* segments in *z*_1:*t*_), counted from the sequence start;
- *B_c,t_*(*j*) = *P*(*x_t_*_+1:*N*_ | *z_t_* = *j*, *c* segments in *z_t_*_:*N*_), counted from the sequence end.

Both counts include the segment that contains *z_t_*. Each lattice follows the standard recursion, except that the segment index remains unchanged during a self-transition and increases by one when the state changes. Partitions with *c* > *K* are never formed. Forward and backward partitions are combined only if the total segment count does not exceed *K*:

- state posterior at *t*: *c*_F_ + *c*_B_ − 1 ≤ *K*, because the two partitions share the segment containing *z_t_*;
- transition *i* → *j* between *t* and *t*+1: *c*_F_ + *c*_B_ − 1 ≤ *K* if *i* = *j* (shared segment), and *c*_F_ + *c*_B_ ≤ *K* if *i* ≠ *j*.

All posteriors are normalized by *P*(**x**, *s_N_* ≤ *K*) = Σ*_c,j_ F_c,N_*(*j*).

In the M-step, the constrained pairwise posteriors are used to re-estimate the transition probabilities. The constrained state posteriors re-estimate the start probabilities and the emission parameters: categorical emission probabilities, or Gaussian means and diagonal variances. The remaining update formulas are unchanged from hmmlearn. After convergence, the most probable path with at most *K* segments is identified using a max-product recursion over the same segment-indexed lattice. This recursion stores back-pointers to (segment, state) pairs and traces back from the best final entry over all *c* ≤ *K*.

Computational cost is O(*NK*²*M*²) time for the pairwise posteriors and O(*NKM*²) for the other passes, with O(*NKM*) memory. We used *K* = 4. Each chromosome arm was fitted using up to 20 random initializations for models with learned emissions (Gaussian state means initialized by k-means with a single initialization) and a single deterministic initialization for models with fixed emissions, retaining the converged model with the highest likelihood. A state never visited during training was retained with a self-transition. Failed initializations were skipped, with fallback to fixed parameters, still under the k-segment constraint, only if all attempts failed. The implementation, including the k-segment Viterbi path, was verified against exhaustive enumeration of all admissible paths in small models (maximum a posteriori path recovered in 300/300 trials). Code is available at https://github.com/recombinationlab/GroupHMM.

The *K* = 4 per chromosome arm allows one segment for an unchanged or wholly homozygous arm, two for terminal LOH, three for interstitial LOH, and four for combinations of these events. Start and transition probabilities were learned independently for each chromosome arm, whereas LOH emission parameters were either fixed or learned according to the model configuration. The fixed-emission models used probabilities of 0.05 and 0.95 for the two LOH states for Gaussian models and 0.1 and 0.9 for Multinomial models, and 0.5 for heterozygosity. Model configurations and parameter settings are summarized in **Tabs.S3-5**.

For comparison, conventional Multinomial and Gaussian HMMs were implemented using the fixed-parameter approach described previously (17). Fixed transition probabilities were scaled by the number of SNP observations or bins per chromosome arm. We also evaluated unconstrained Multinomial HMMs with learned parameters.

### Adaptive SNP binning

For Gaussian HMMs, we evaluated fixed bins containing 10 or 40 phased SNPs and adaptive bins containing 10-40 SNPs. Adaptive bin size was determined from either per-cell sequencing coverage or the number of informative phased SNPs, using chromosome 19, the smallest mouse chromosome, to calibrate SNP-based adaptation and avoid excessive binning in sparsely sampled chromosomes. The adaptive schemes were evaluated with both fixed-transition and k-segment Gaussian HMMs (**Tab.S4**).

### LOH filtering and segment merging

HMM segments were classified as LOH based on observed haplotype fractions or learned emission probabilities, depending on the model configuration. LOH calls required a haplotype fraction or emission probability meeting the model-specific threshold, a minimum length of 10 Mb, and at least 20 informative phased SNPs. Gaussian HMM segments were refined to SNP-level boundaries using the likelihood-ratio procedure described previously (17). Filtered LOH segments belonging to the same haplotype were merged when adjacent or separated by a non-LOH gap shorter than 10 Mb. Merging was performed iteratively within each chromosome arm, without extending LOH into terminal non-LOH regions.

### LOH benchmarking

HMM performance was evaluated against the sci-L3-Strand-seq reference comprising 2,395 LOH and 22,012 non-LOH segments from 931 Patski cells. Reference LOH included segmental deletions, monosomy, segmental cnLOH, and UPD. An HMM call was considered a TP when it overlapped at least 60% of a reference LOH segment. FP calls were counted when filtered HMM LOH regions overlapped reference non-LOH segments, without a minimum overlap threshold.

Recall was calculated as TP/2,395, precision as TP/(TP+FP), and FPR as FP/22,012. Model performance was compared using precision–recall and receiver operating characteristic (ROC) plots. The distance from ideal precision and recall was calculated as

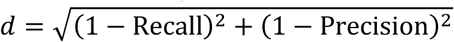

Whole-chromosome and segmental LOH were analyzed separately; HMM calls spanning more than 90% of a chromosome were classified as whole-chromosome events for FP categorization.

### Haplotype-aware validation of apparent FP calls

To evaluate apparent FP calls, sci-L3-Strand-seq reads were separated by parental haplotype as described previously. Haplotype-specific strand-orientation changes and read coverage were examined to distinguish genuine LOH from copy-number gains, drop-to-zero (DTZ) regions, or unsupported HMM segmentation.

For each of three k-segment HMM configurations, 30 stand-alone and 30 LOH-adjacent apparent FP calls were manually examined. LOH-adjacent FPs were defined as HMM calls extending no more than 10 Mb beyond a reference LOH boundary. Calls were classified as likely genuine LOH, DTZ-associated FP, gain-associated FP, or FP lacking sufficient haplotype-aware evidence (**Tab.2, S6-11**).

### Clonal analysis of apparent FP calls

LOH profiles were encoded as binary vectors at 1-Mb resolution for each cell and parental haplotype. Hierarchical clustering was performed using `sklearn`, with 20 clusters, initially using genome-wide apparent FP calls and subsequently chromosome-specific LOH profiles. Recurrent apparent FP calls were evaluated against reference TP events and haplotype-aware evidence to identify likely genuine LOH shared among cells (**Fig.7, S12-13; Tab.S12-14**).

## Supplemental Figures

**Figure S1.**
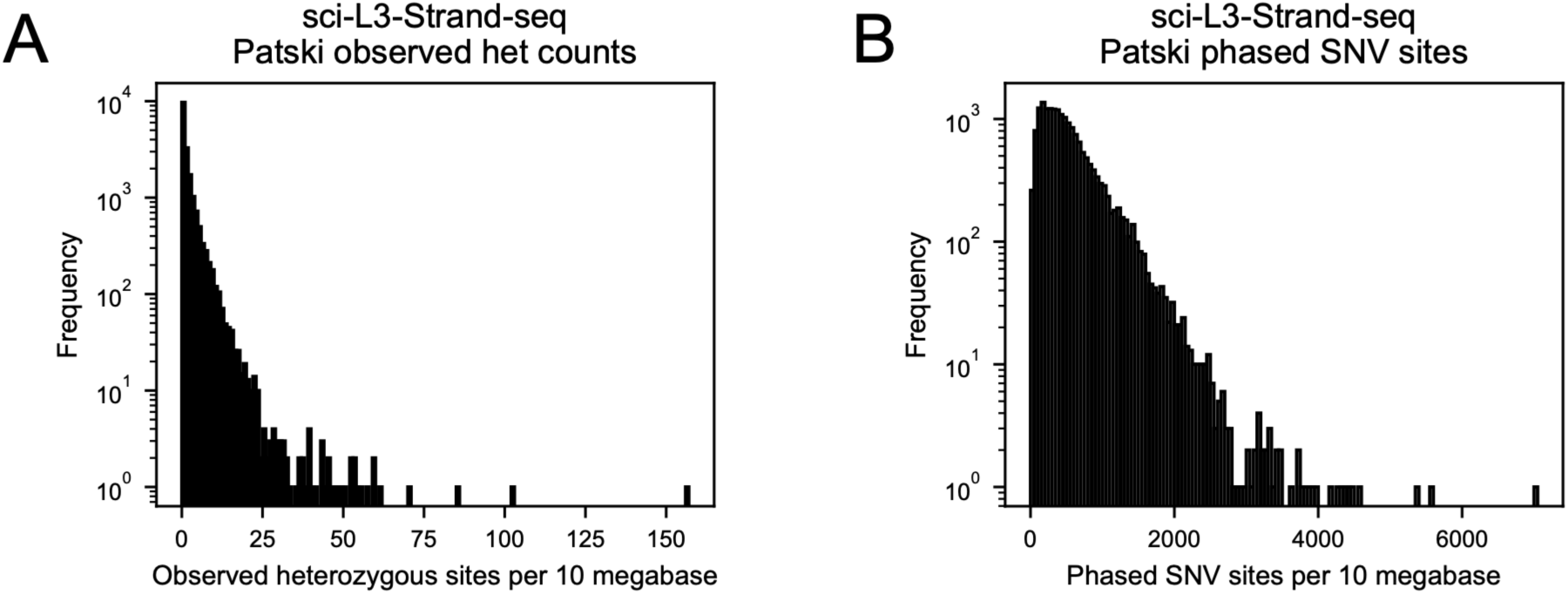
Sparse single-cell sequencing rarely captures both alleles at heterozygous sites. Histograms show the distributions of directly observed heterozygous sites (A) and informative phased SNVs (B) per 10 Mb across 18,620 chromosomes from 931 Patski cells (logarithmic y-axes). **A.** Directly observed heterozygosity is sparse, with a median of 0.93 heterozygous sites per 10 Mb. Only 629 chromosomes (3.4%) contain ≥10 observed heterozygous sites per 10 Mb, and just 15 (0.08%) contain ≥50. **B.** In contrast, phased SNVs are substantially more abundant, with a mode of approximately 150-200 and a median of approximately 440 sites per 10 Mb. This increased availability of informative haplotype observations enables LOH inference from sparse single-cell WGS despite the rarity of directly observed heterozygosity.

**Figure S2.**
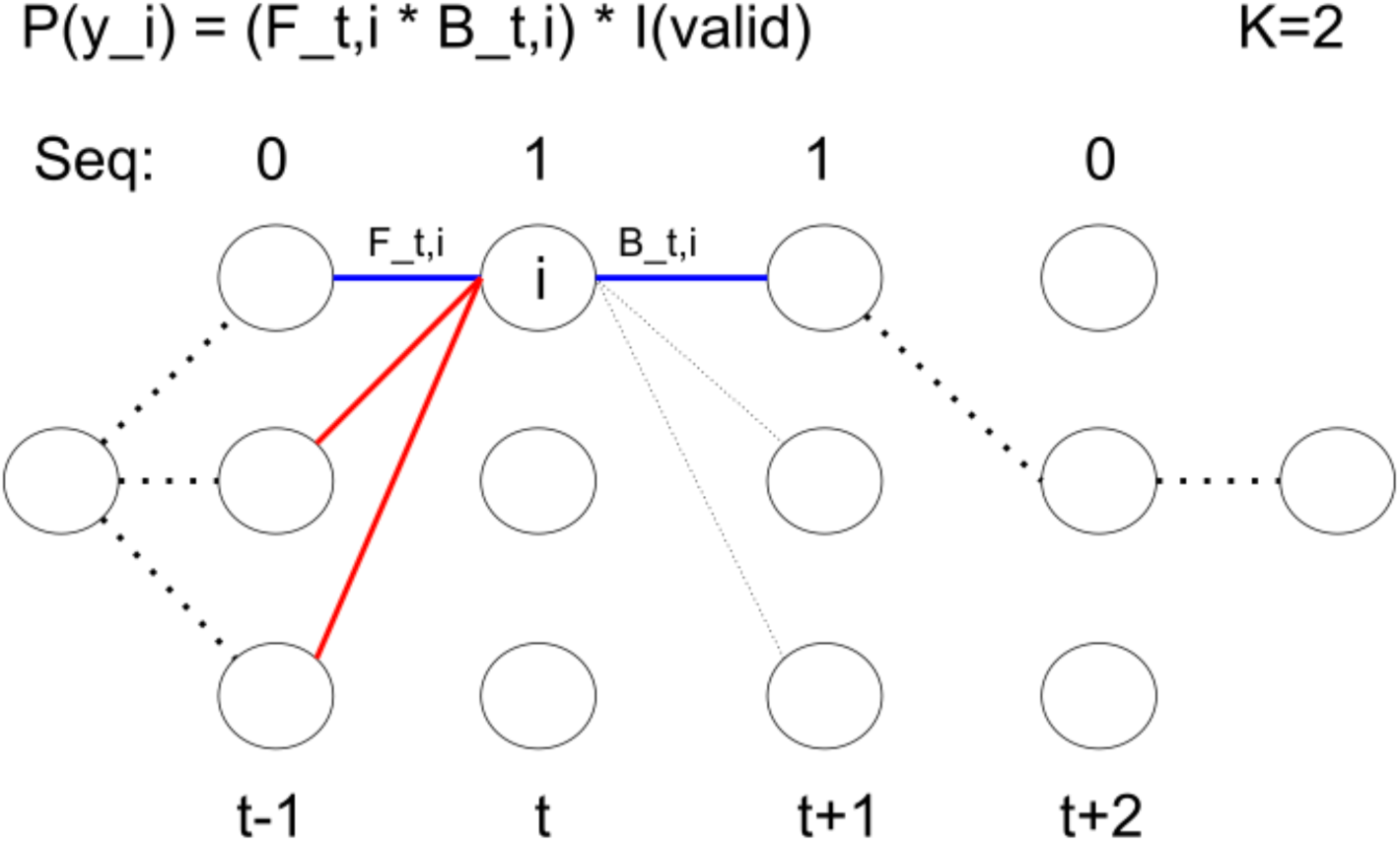
Segment-count compatibility in the *k*-segment constrained HMM. Schematic of a three-state HMM with four observations (0, 1, 1, 0), illustrating constrained forward–backward inference with a maximum of *K* = 2 segments. Nodes represent hidden states at successive time steps. The blue edges indicate compatible forward and backward partitions: the forward path contains one segment, while the backward path contains two, sharing the segment at time *t* and therefore satisfying the two-segment constraint. In contrast, the red edges introduce an additional state transition between *t* − 1 and *t*, producing three segments when combined with the backward path and thus violating the constraint. Only compatible partitions contribute to the constrained state posterior used for parameter learning.

**Figure S3.**
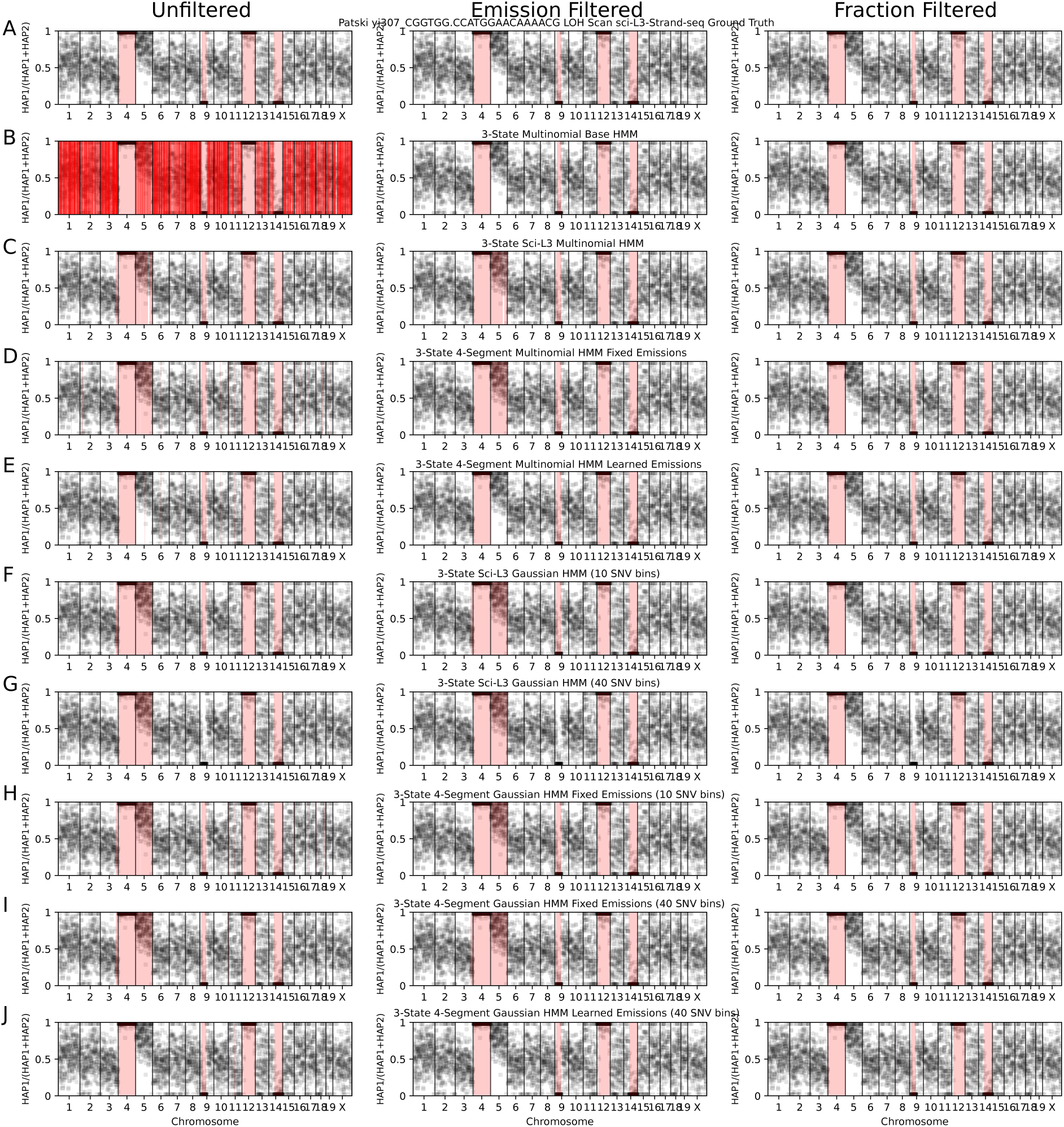
Effects of LOH filtering across HMM configurations in a median-coverage Patski cell. LOH calls for Patski cell yi307_CGGTGG.CCATGGAACAAAACG (0.53% coverage), also shown in Fig.3, before filtering (left), after emission-probability filtering (middle), and after observed haplotype-fraction filtering (right). Points represent haplotype fractions calculated from 40-SNP bins and plotted at bin midpoints; red shading indicates LOH calls. The sci-L3-Strand-seq reference identifies LOH on chromosomes 4, 9, 12, and 14. The comparison illustrates how filtering reduces spurious LOH calls, while the *k*-segment constraint controls over-segmentation during parameter learning. **A.** sci-L3-Strand-seq reference LOH calls. **B.** Standard three-state Multinomial HMM with fixed emissions and unconstrained learned transitions. The model produces 9,944 chromosome segments before filtering, reflecting severe over-segmentation. Filtering removes most short LOH calls but does not fully recover the reference events. **C.** Three-state sci-L3 Multinomial HMM with fixed emissions and transitions, recovering all four reference LOH events without FP calls after filtering. **D,E.** Four-segment Multinomial HMMs with fixed (D) or learned (E) emissions and learned transitions. The *k*-segment constraint suppresses excessive segmentation while retaining the reference LOH events. **F,G.** Three-state sci-L3 Gaussian HMMs with fixed parameters and 10-SNP (F) or 40-SNP (G) bins, illustrating the effects of bin size on segmental LOH detection. **H,I.** Four-segment Gaussian HMMs with fixed emissions, learned transitions, and 10-SNP (H) or 40-SNP (I) bins, illustrating improved segmental LOH detection with constrained transition learning. **J.** Four-segment Gaussian HMM with learned emissions and transitions using 40-SNP bins.

**Figure S4.**
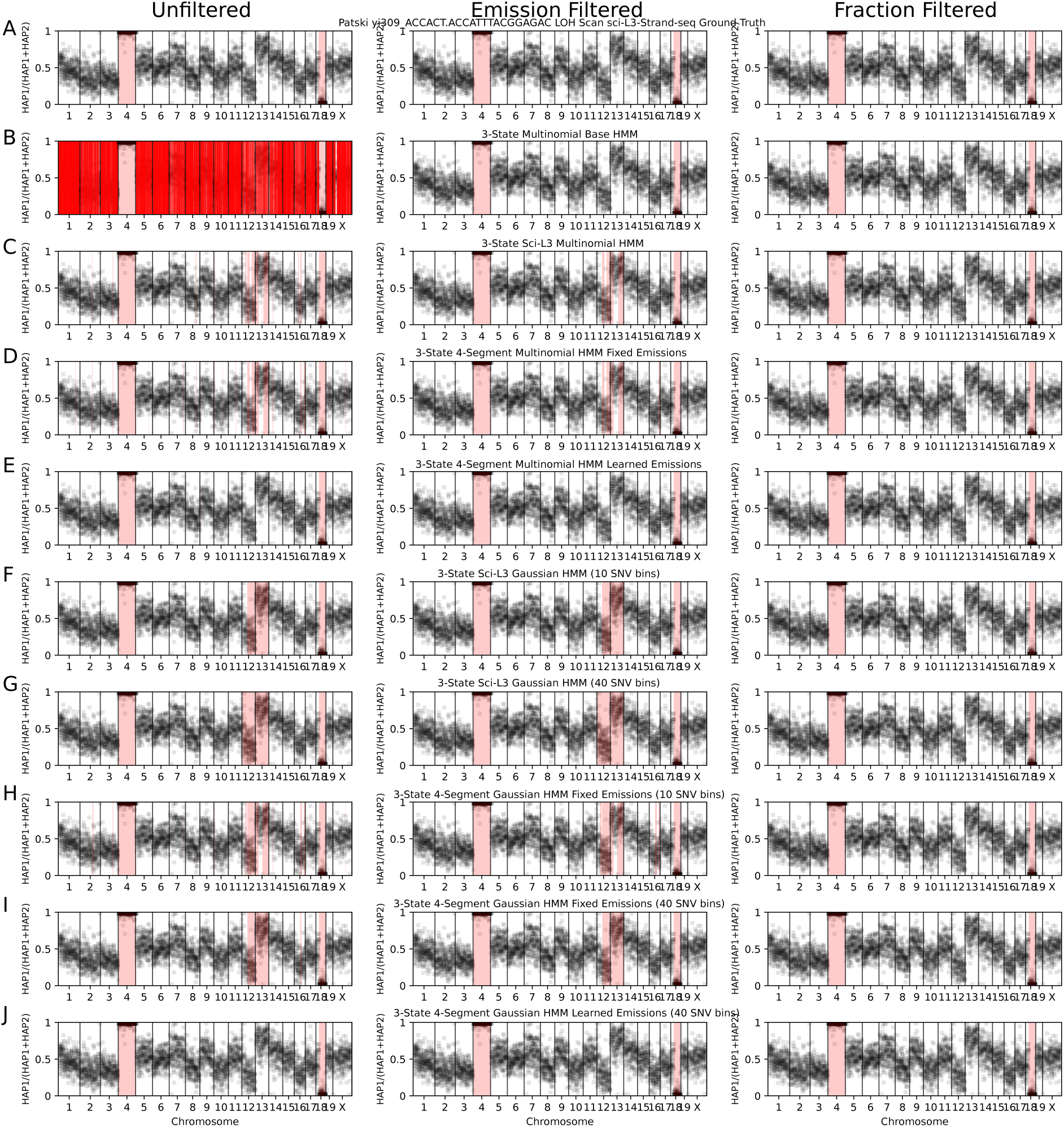
Effects of LOH filtering across HMM configurations in a higher-coverage Patski cell. LOH predictions for Patski cell yi309_ACCACT.ACCATTTACGGAGAC (1.43% coverage). Panel organization, HMM configurations (A–J), filtering strategies, and color conventions are identical to **Fig.S3**. The sci-L3-Strand-seq reference identifies whole-chromosome LOH on chromosome 4 and segmental LOH on chromosome 18. The comparison illustrates the effects of HMM configuration and filtering on LOH detection at higher sequencing coverage.

**Figure S5.**
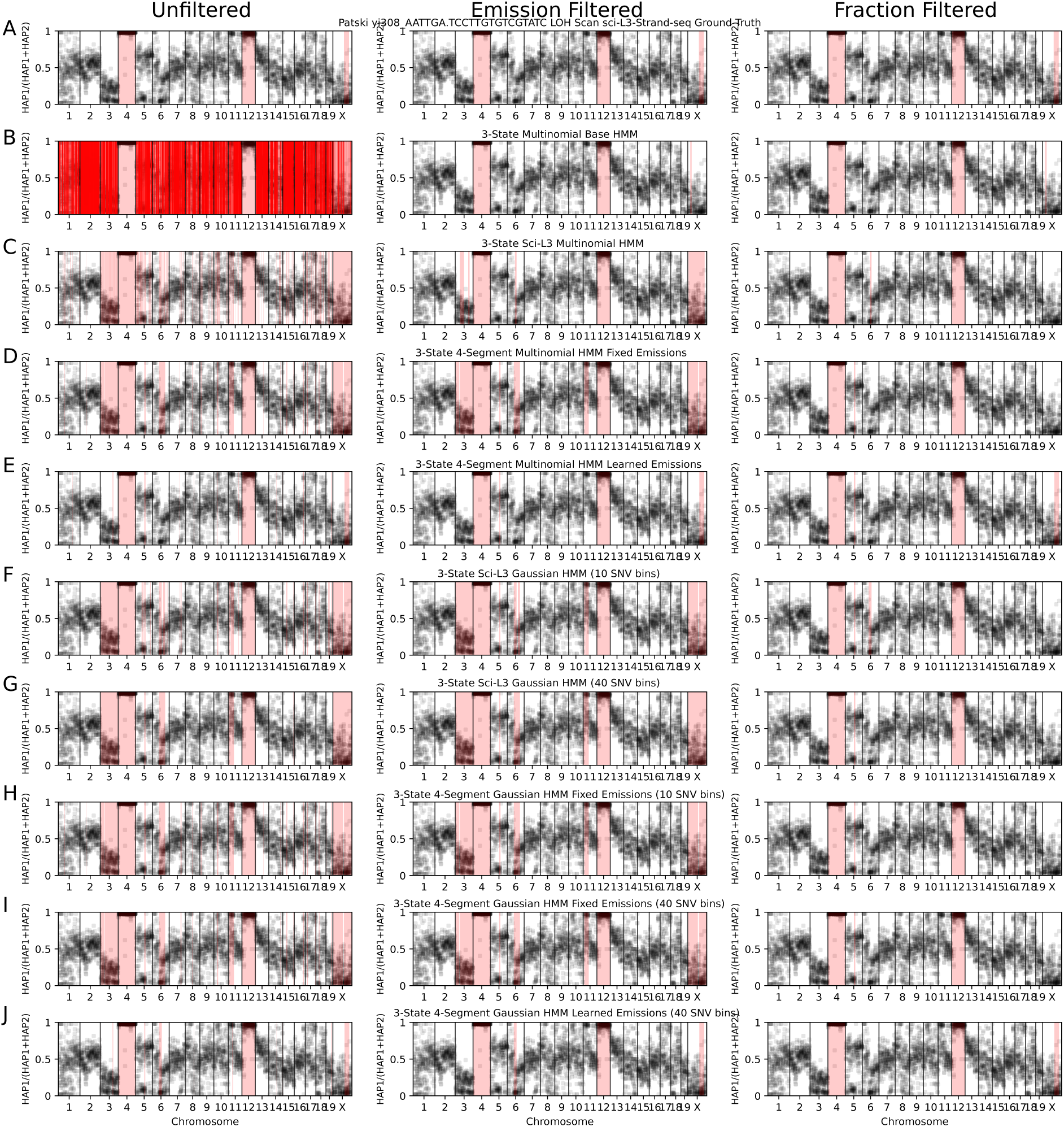
Effects of LOH filtering across HMM configurations in a high-coverage Patski cell. LOH predictions for Patski cell yi308_AATTGA.TCCTTGTGTCGTATC (3.16% coverage). Panel organization, HMM configurations (A–J), filtering strategies, and color conventions are identical to **Fig.S3**. The sci-L3-Strand-seq reference identifies whole-chromosome LOH on chromosomes 4 and 12 and segmental LOH on chromosome X. The comparison illustrates how the *k*-segment constraint limits over-segmentation at high sequencing coverage, particularly relative to the standard and fixed-transition Multinomial HMMs.

**Figure S6.**
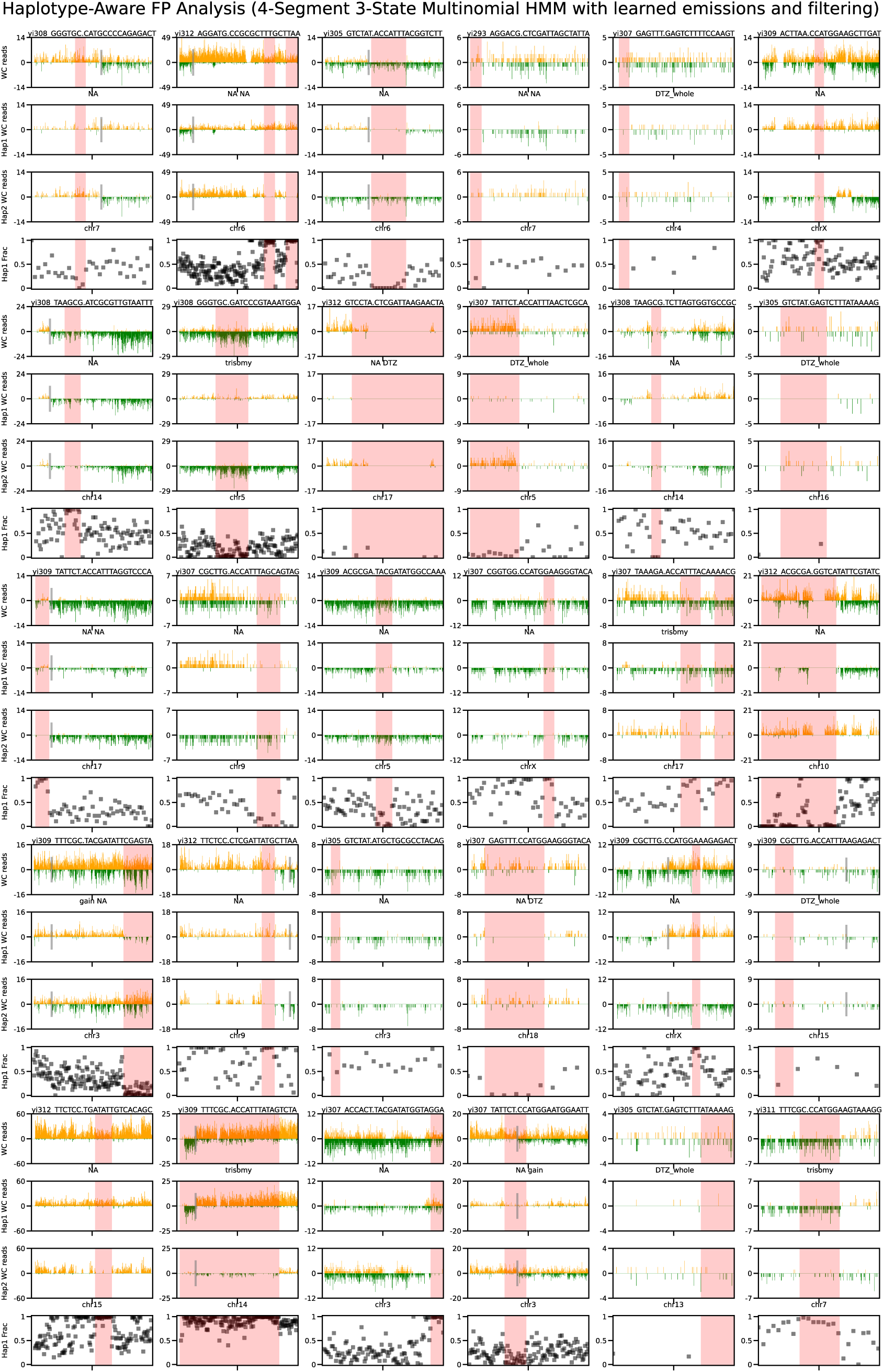
Haplotype-aware analysis of stand-alone apparent FP LOH calls from the *k*-segment Multinomial HMM with learned emissions. Haplotype-aware sci-L3-Strand-seq analysis of 30 stand-alone apparent FP LOH calls filtered by learned emission probabilities. Each example shows strand-specific reads, haplotype 1 and haplotype 2 reads, and haplotype fractions. Red shading indicates HMM LOH calls, yellow indicates sci-L3-Strand-seq reference LOH, and overlapping regions appear orange. Manual classifications are provided in **Tab.S6**.

**Figure S7.**
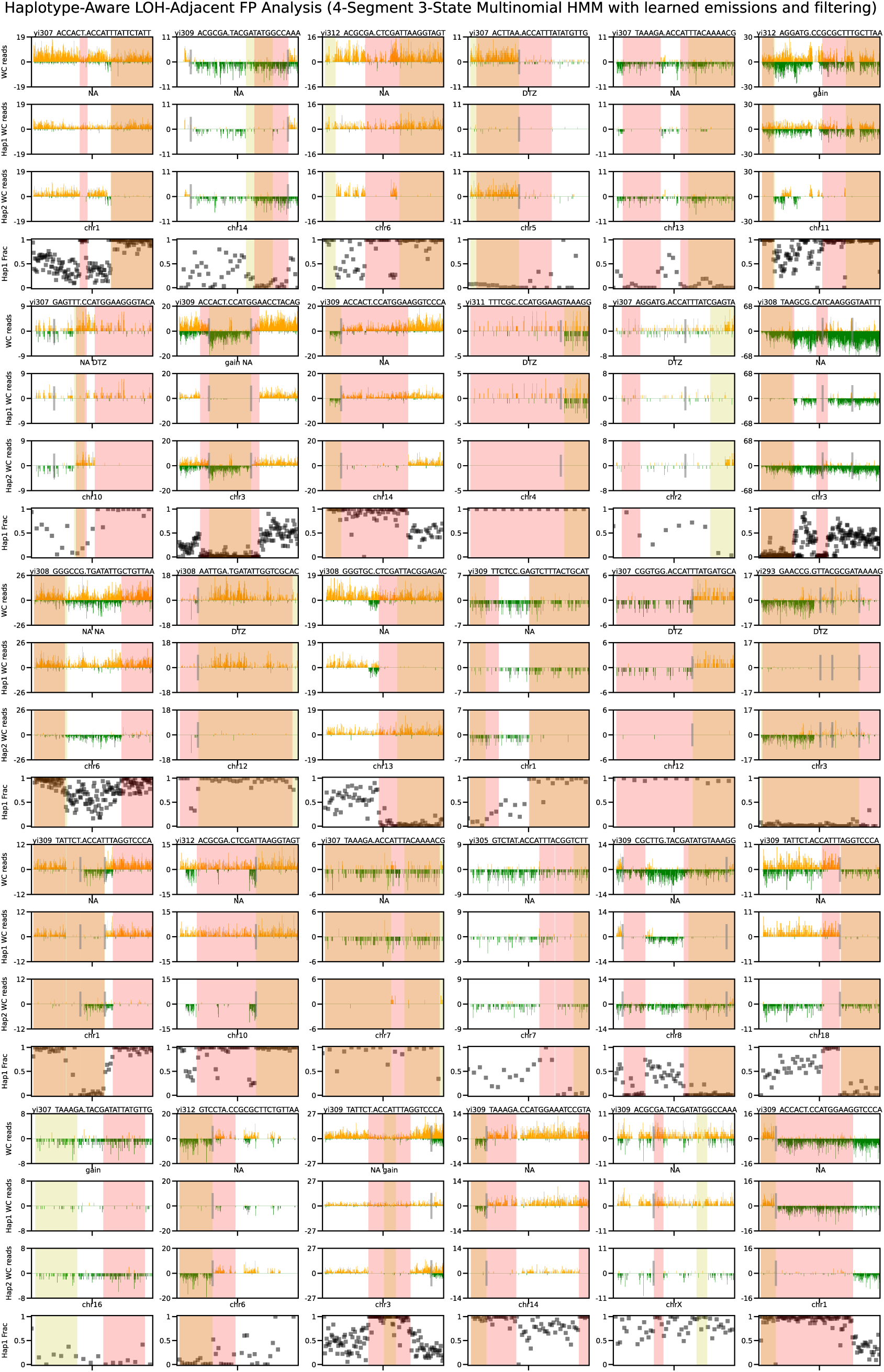
Haplotype-aware analysis of LOH-adjacent apparent FP calls from the *k*-segment Multinomial HMM with learned emissions. Analysis of 30 LOH-adjacent apparent FP calls using the same HMM configuration and plotting conventions as **Fig.S6**. These calls overlap or extend beyond sci-L3-Strand-seq reference LOH boundaries. Manual classifications are provided in **Tab.S7**.

**Figure S8.**
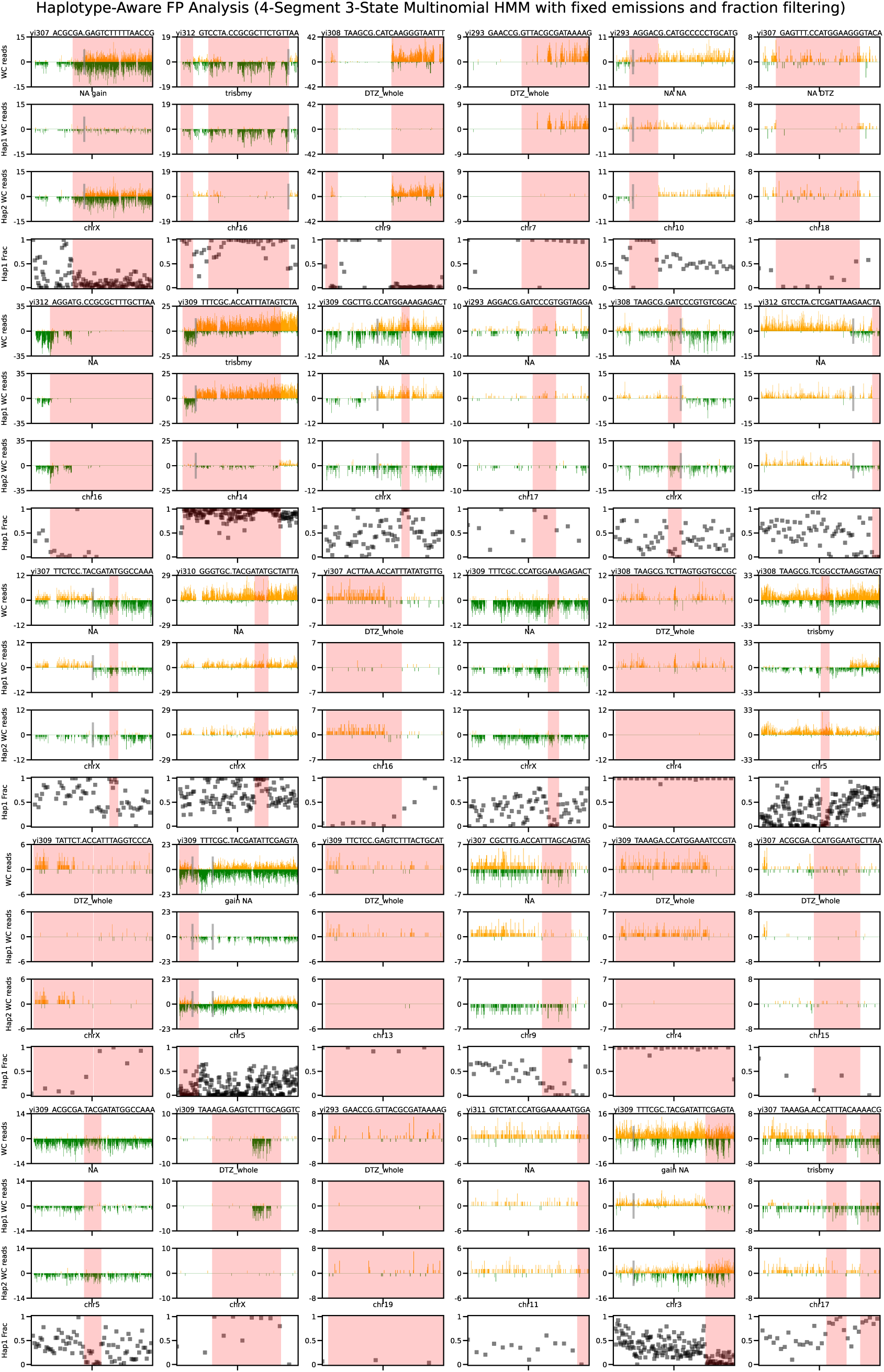
Haplotype-aware analysis of stand-alone apparent FP LOH calls from the *k*-segment Multinomial HMM with fixed emissions. Analysis of 30 stand-alone apparent FP LOH calls from the fixed-emission Multinomial HMM with haplotype-fraction filtering. Plotting conventions are as in **Fig.S6**. Manual classifications are provided in **Tab.S8**.

**Figure S9.**
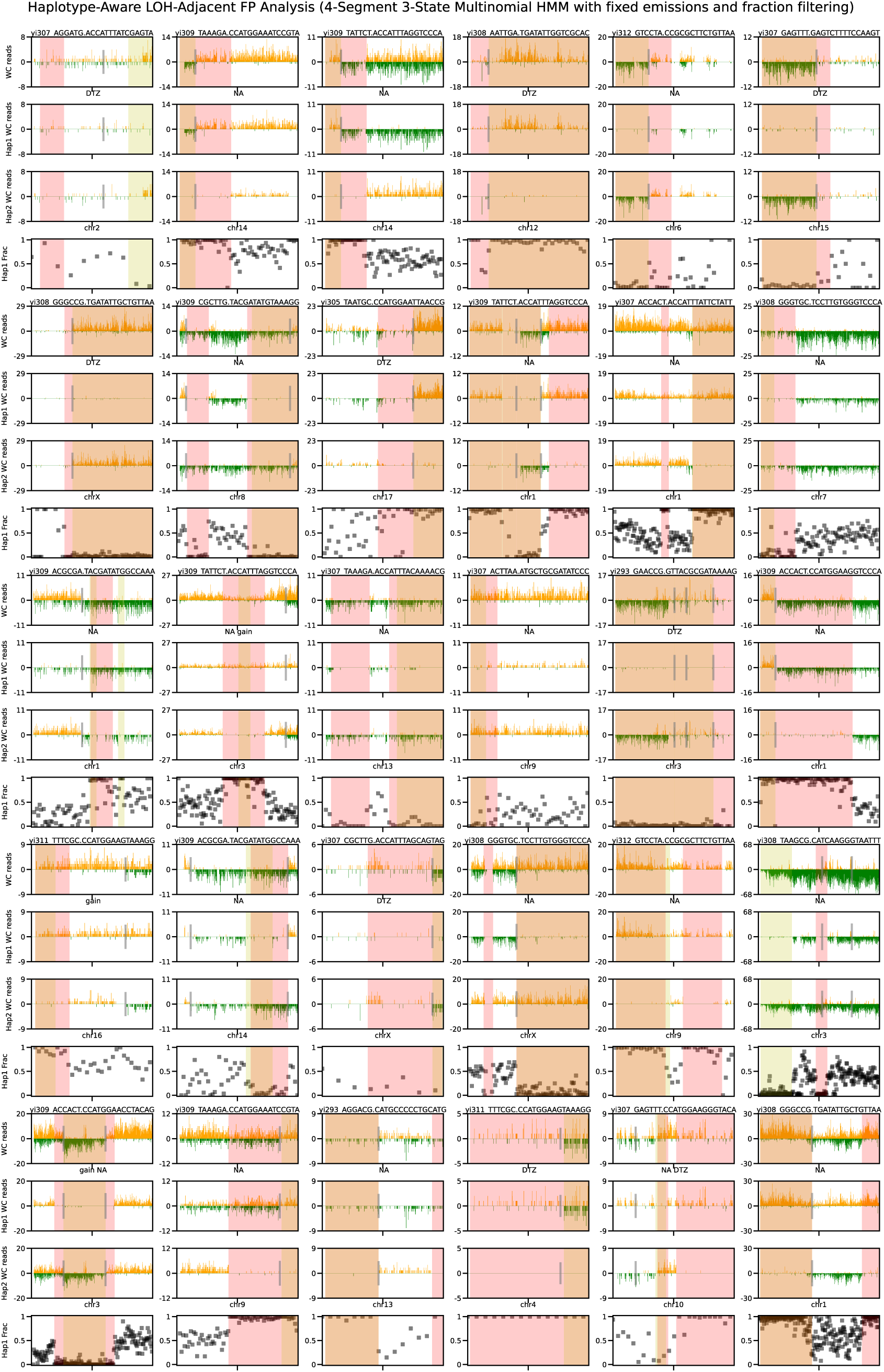
Haplotype-aware analysis of LOH-adjacent apparent FP calls from the *k*-segment Multinomial HMM with fixed emissions. Analysis of 30 LOH-adjacent apparent FP calls from the fixed-emission Multinomial HMM with haplotype-fraction filtering. Plotting conventions are as in **Fig.S6**. Manual classifications are provided in **Tab.S9**.

**Figure S10.**
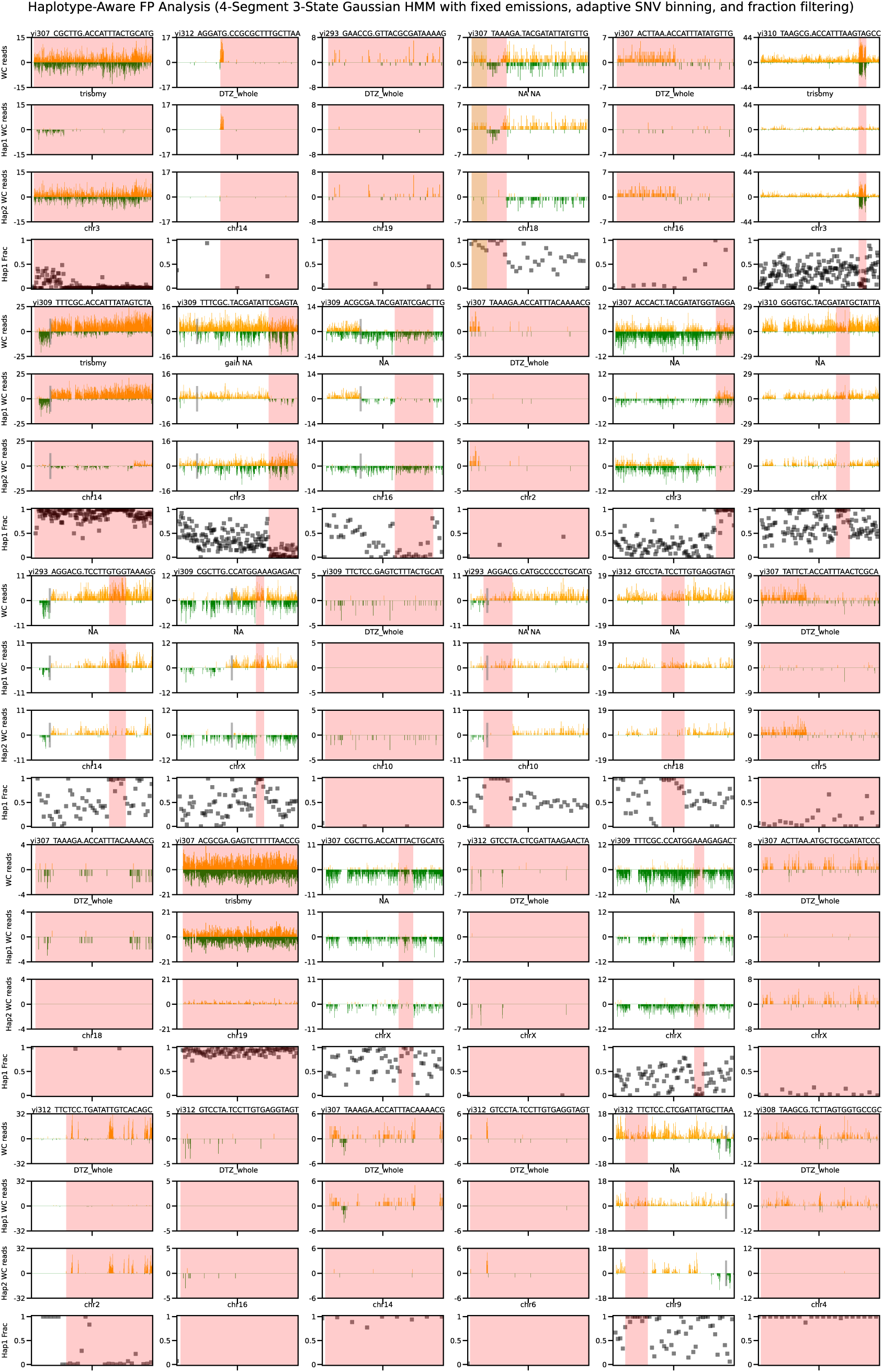
Haplotype-aware analysis of stand-alone apparent FP LOH calls from the *k*-segment Gaussian HMM with adaptive SNP binning. Analysis of 30 stand-alone apparent FP LOH calls from the Gaussian HMM with fixed emissions, adaptive SNP binning, and haplotype-fraction filtering. Plotting conventions are as in **Fig.S6**. Manual classifications are provided in **Tab.S10**.

**Figure S11.**
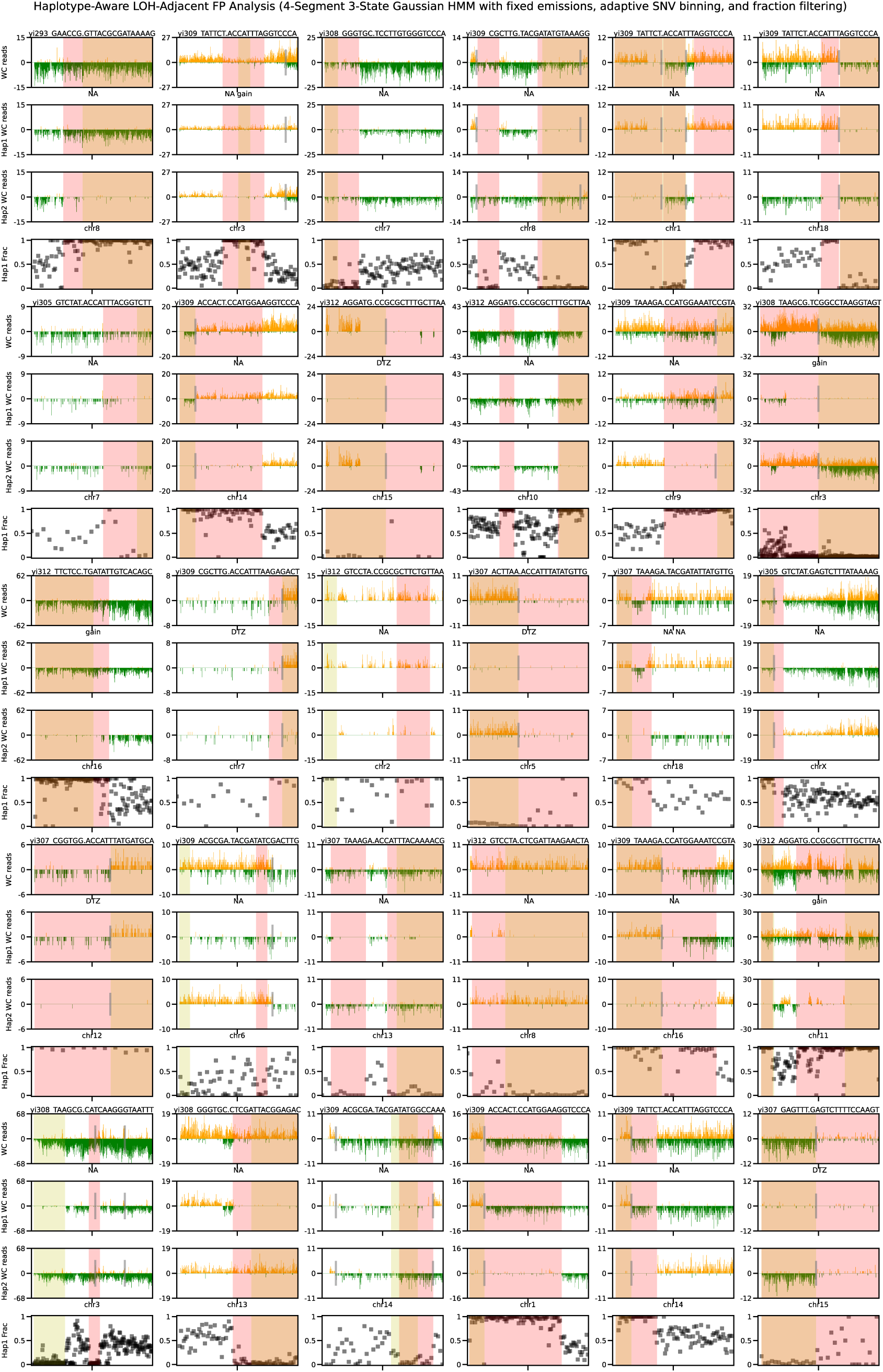
Haplotype-aware analysis of LOH-adjacent apparent FP calls from the *k*-segment Gaussian HMM with adaptive SNP binning. Analysis of 30 LOH-adjacent apparent FP calls from the Gaussian HMM with fixed emissions, adaptive SNP binning, and haplotype-fraction filtering. Plotting conventions are as in **Fig.S6**. Manual classifications are provided in **Tab.S11**.

**Figure S12.**
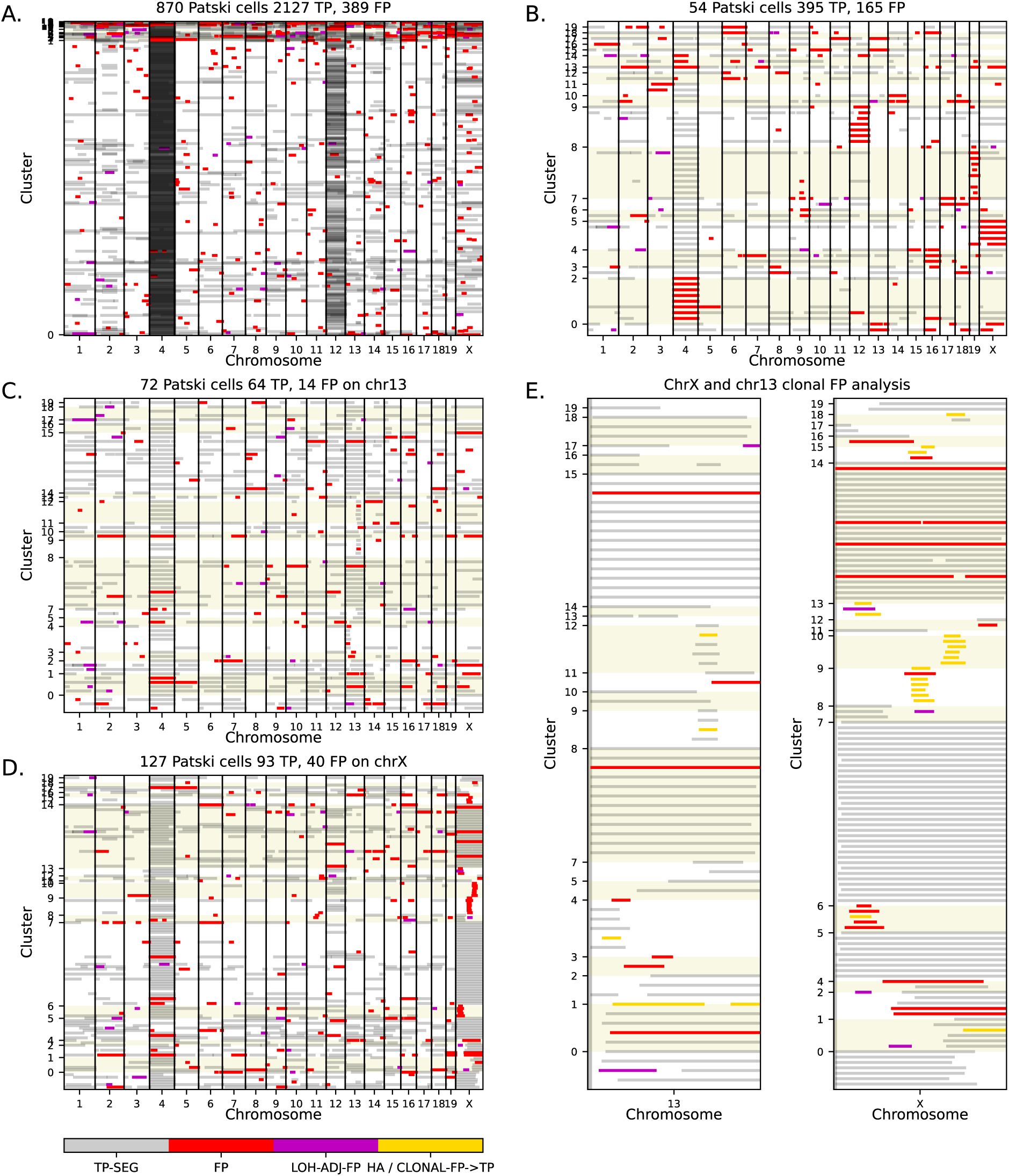
Clonal analysis of apparent FP LOH calls from the *k*-segment Multinomial HMM with fixed emissions. Hierarchical clustering of LOH profiles from the four-segment, three-state Multinomial HMM with fixed emissions and haplotype-fraction filtering. Panel organization, color conventions, and clustering procedures are as described in Fig.7. Recurrent apparent FP calls on chr13 and chrX were evaluated using haplotype-aware analysis, with manual classifications provided in **Tab.S13**.

**Figure S13.**
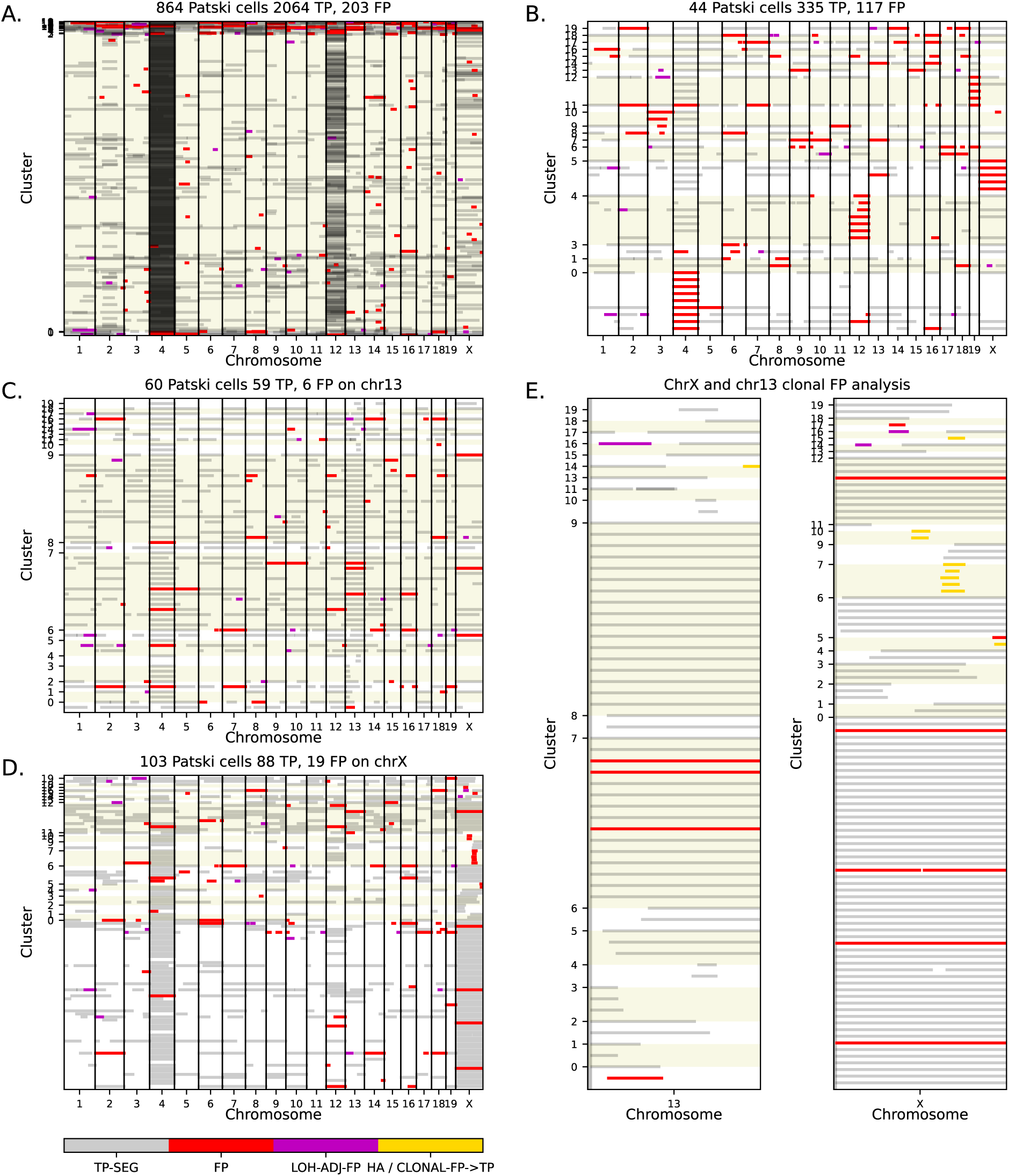
Clonal analysis of apparent FP LOH calls from the *k*-segment Gaussian HMM with adaptive SNP binning. Hierarchical clustering of LOH profiles from the four-segment, three-state Gaussian HMM with fixed emissions, adaptive SNP binning, and haplotype-fraction filtering. Panel organization, color conventions, and clustering procedures are as described in Fig.7. Recurrent apparent FP calls on chr13 and chrX were evaluated using haplotype-aware analysis, with manual classifications provided in **Tab.S14**.

## Notes

### Competing Interest Statement

The authors have declared no competing interest.

